# Structural and mechanistic insights into *Streptococcus pneumoniae* NADPH oxidase

**DOI:** 10.1101/2023.10.17.562464

**Authors:** Victor R. A. Dubach, Pablo San Segundo-Acosta, Bonnie J. Murphy

## Abstract

NADPH oxidases (NOXs) play a major role in the physiology of eukaryotic cells by mediating the production of reactive oxygen species (ROS). Evolutionarily distant proteins sharing the NOX catalytic core have been recently described in Bacteria. Among them, the *Streptococcus pneumonia*e NOX (SpNOX) has been proposed as a model for the study of NOXs due to its high activity and stability in detergent micelles. Here, we report high-resolution cryo-EM structures of substrate-free and stably reduced NADH-bound SpNOX, and of the NADPH-bound SpNOX and a Phe397Ala mutant under turnover conditions. In combination with structure-guided mutagenesis and biochemical analyses, we provide the structural basis for constitutive activity, the lack of substrate specificity towards NADPH and the electron transfer pathway. Additionally, we shed light on the catalytic regulation by the C-terminal tail residue Phe397 and the potential *in vivo* function of this protein.

## Introduction

Reactive oxygen species (ROS), such as superoxide anion radical (O_2_^•–^) and hydrogen peroxide (H_2_O_2_), are highly reactive chemical species that play essential roles in immunity, cell signaling, ageing and cancer [1]. The NADPH oxidase (NOX) family of enzymes (E.C. 1.6.3.1) generate ROS as their primary product and therefore play a major role in ROS homeostasis. NOXs belong to the ferric reductase transmembrane component-like domain (FRD) superfamily, and are composed of a catalytic core with two conserved domains: a domain with six transmembrane helices (TM domain) coordinating two hemes via four conserved histidine residues; and a C-terminal cytosolic dehydrogenase domain (DH domain) containing the FAD- and NADPH-binding sites (Fig. 1a). The DH domain catalyzes a hydride transfer to FAD, which subsequently transfers electrons stepwise to the two transmembrane hemes. The outer heme reduces oxygen to generate O_2_^•–^ or H_2_O_2_ [2, 3].

**Fig. 1.**
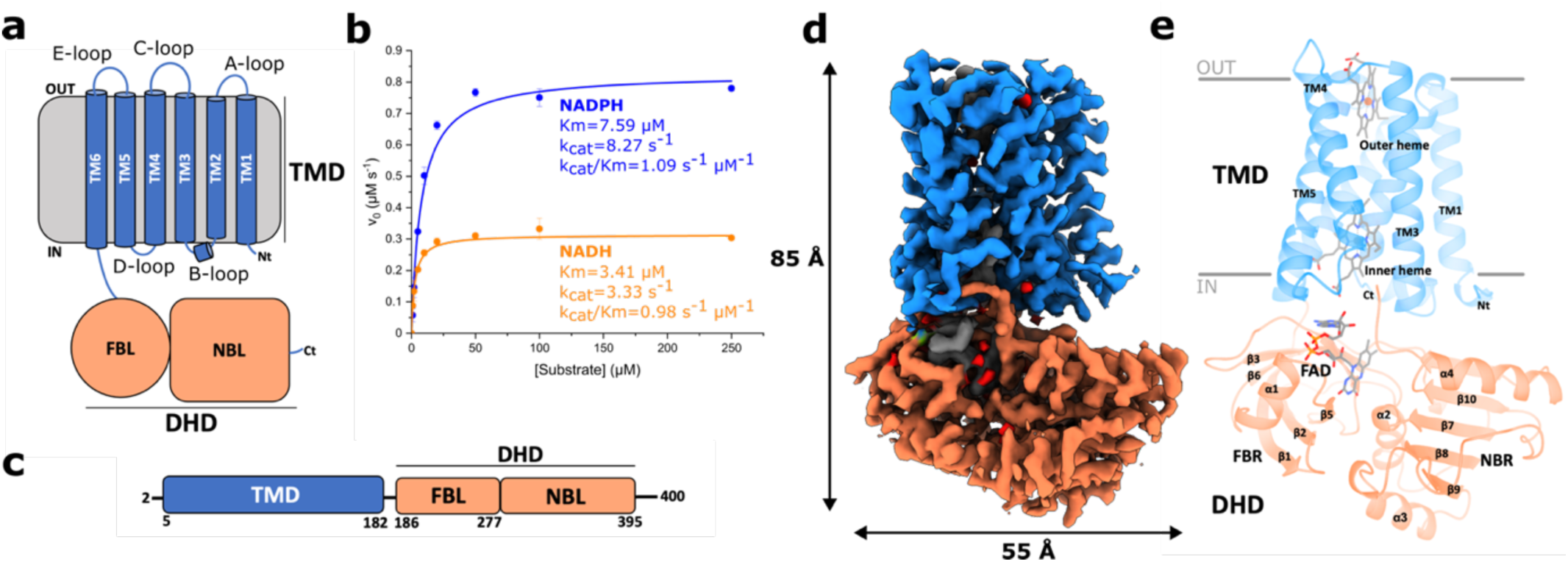
SpNOX displays the canonical NOX structure and NAD(P)H-oxidase activity. **a,** SpNOX is composed of a transmembrane domain (TMD) and a dehydrogenase domain (DHD) subdivided into an FAD-binding lobe (FBL) and an NAD(P)H-binding lobe (NBL). **b,** The activity of SpNOX under steady state conditions was measured using a cytochrome c reduction assay. Apparent K_m_ and k_cat_ values were obtained by fitting the data to the Michaelis-Menten equation. Mean values of three technical replicates are plotted and the SEM is indicated. Data for individual replicates are available in the source data. **c**, Schematic view of SpNOX domain boundaries**. d,** Side view of the cryo-EM map of substrate-free SpNOX. TMD and DHD are colored blue and coral, respectively. The same color pattern is used throughout the article unless otherwise stated. **e,** Structure of substrate-free SpNOX in cartoon representation. Membrane boundaries are represented as grey lines.

Although NOXs were first thought to be exclusive to eukaryotic genomes, genes encoding proteins sharing the same catalytic core have more recently been found in Bacteria [4, 5]. With the exception of cyanobacterial NOX5, which seems to have arisen by horizontal gene transfer from eukaryotic donors, bacterial NOX homologues are evolutionarily distant from eukaryotic NOXs and consist of a single polypeptide chain with DH and TM domains [5]. The best characterized member is the *Streptococcus pneumoniae* NOX (SpNOX). SpNOX is a 46-kDa protein that can be produced with correct heme incorporation in high yields, and shows high stability and robust activity when solubilized in lauryl maltose neopentyl glycol (LMNG) [4, 6].

Structural studies of NOX proteins are generally difficult due to low yields after recombinant expression, loss of cofactors during purification and flexibility. The first structure of a NOX protein was the crystal structure of *Cylindrospermum stagnale* NOX5 (csNOX5) separate TM and DH domains, which gave insight into the catalytic core and the oxygen-reduction center [7]. However, despite significant efforts, obtaining full-length NOX crystals diffracting to high resolution has proved to be challenging [6, 7]. Nevertheless, the use of cryo-EM has allowed to solve the structures of mouse and human dual oxidases (DUOX1) [8, 9], and the inactive human core NOX2 complex [10, 11]. These structures have provided essential information about subunit interactions and catalytic regulation of eukaryotic NOXs, but large questions remain regarding the mechanism of activation and catalytic turnover. Moreover, no structure of a NOX-like bacterial protein is currently available. Due to the relatively simple expression and purification of SpNOX, and its homology to eukaryotic NOX proteins, a structural characterization of SpNOX would be valuable for its application as a model system of NADPH oxidases.

Here, we present high-resolution cryo-EM structures of SpNOX obtained in substrate-free conditions, under stably reducing conditions with NADH and under turnover conditions with NADPH. Compared to csNOX5, DUOX1 and NOX2, SpNOX is smaller, has shorter extracellular loops and, like the mammalian NOX4, it is constitutively active. These are the first structures of a bacterial NOX-like protein and, together with structure-based mutagenesis, they provide insights into the electron transfer pathway, catalytic mechanism and constitutive activity.

## Results

### SpNOX purification, analysis and structure determination

*S. pneumoniae* NOX was expressed in *Escherichia coli* and purified as previously described with minor modifications (see Methods). The protein was solubilized using LMNG and showed a single band in SDS-PAGE and a homogeneous single peak on SEC, corresponding to the monomeric protein (Supplementary Fig. 1a,b). Quick removal of imidazole after Ni-NTA purification was crucial to avoid protein aggregation. The UV-vis spectrum of the purified protein showed the characteristic Soret peak at 414 nm, indicating correct heme incorporation (Supplementary Fig. 1c). Oxidation of NADPH could be directly followed monitoring absorbance at 340 nm under aerobic conditions. A similar experiment in the absence of O_2_ showed only initial activity due to residual O_2_ in the buffer and/or reduction of SpNOX cofactors before the oxidation activity ceased, confirming that O_2_ can act as an electron acceptor for SpNOX (Supplementary Fig. 1d). NADPH oxidation was also measured with a cytochrome c reduction assay [4] (Fig. 1b), and Michaelis-Menten analysis gave an apparent K_m_ value of 7.59 µM and an apparent k_cat_ of 8.27 s^-1^. The protein also exhibited NADH-driven oxygen reduction, with an apparent K_m_ value of 3.41 µM, and an apparent k_cat_ of 3.33 s^-1^. Cytochrome c reduction assays under anaerobic conditions revealed direct electron transfer to cytochrome c (Supplementary Fig. 1e), likely from the outer heme, and to superoxide dismutase (SOD) (Supplementary Fig. 1f). Therefore, the rate of cytochrome c reduction by the anion superoxide could not be estimated with SOD inhibition. Our results, obtained under both aerobic and anaerobic conditions, indicate that previous efforts to measure specific superoxide production using SOD inhibition may have been subject to unforeseen artifacts. The initial rates of cytochrome c reduction under aerobic and anaerobic conditions were very similar (Fig. 1g), suggesting cytochrome c as the major direct electron acceptor in the *in vitro* assay.

Cryo-EM single-particle analysis of SpNOX was performed in LMNG micelles, and showed that the protein preparation was homogeneous (Methods and Supplementary Figs. 2-5). In addition to a structure of SpNOX in the absence of electron donor, we also determined structures of the stably reduced protein bound to NADH under anaerobic conditions, and of the WT protein and a Phe397Ala mutant under turnover with NADPH and O_2_ present. Despite the relatively small size of the protein (46 kDa) for single-particle analysis, the final maps reached nominal resolutions ranging from 2.2 to 2.6 Å (Table 1 and Supplementary Figs. 6,7), allowing us to resolve essentially all residues. The TM and DH domains reached similar overall local resolutions, with only small regions of the NAD(P)H-binding region being less well resolved (Supplementary Fig. 6). In contrast to previously published small-angle neutron scattering (SANS) data indicating potential interdomain flexibility [6], our results indicate that the DH domain interacts rigidly with the TM domain, thus facilitating particle alignment.

**Table 1.** Cryo-EM data collection, refinement and validation statistics.

|  | #1 Substrate-free<br>(EMDB-18644)<br>(PDB 8QT6) | #2 NADPH-bound<br>(EMDB-18645)<br>(PDB 8QT7) | #3 NADH-bound<br>(EMDB-18646)<br>(PDB 8QT9) | #4 F397A<br>(EMDB-18647)<br>(PDB 8QTA) |
| --- | --- | --- | --- | --- |
| **Data collection and processing** |  |  |  |  |
| Magnification | 215,000 x | 215,000 x | 215,000 x | 215,000x |
| Voltage (kV) | 300 | 300 | 300 | 300 |
| Electron exposure (e<sup>-</sup>/Å<sup>2</sup>) | 70 | 70 | 70 | 70 |
| Defocus range (μm) | -0.7 to -2.1 | -0.7 to -2.1 | -0.7 to -2.1 | -0.7 to -2.1 |
| Pixel size (Å) | 0.573 | 0.573 | 0.573 | 0.573 |
| Symmetry imposed | C1 | C1 | C1 | C1 |
| Initial particle images (no.) | 3,376,347 (Topaz) | 6,641,288 (Topaz) | 5,571,895 (Topaz) | 5,003,252 (Topaz) |
|  | 3,115,826 (Topaz) | 4,948,429 (Topaz) | 5,183,375 (Topaz) | 4,668,309 (Topaz) |
|  | 2,273,739<br>(crYOLO) | 2,868,991 (crYOLO) | 3,583,885<br>(crYOLO) | 4,145,972 (crYOLO) |
| Final particle images (no.) | 397,972 | 591,137 | 697,211 | 546,234 |
| Map resolution (Å) | 2.29 | 2.20 | 2.36 | 2.64 |
| FSC threshold | 0.143 | 0.143 | 0.143 | 0.143 |
| Map resolution range (Å) | 2.25-26.65 | 2.05-32.20 | 2.11-36.46 | 2.50-9.30 |
| **Refinement** |  |  |  |  |
| Initial model used (PDB code) | AlphaAFold<br>model (AF-Q8CZ28) | Substrate-free<br>model | Substrate-free<br>model | NADPH-bound<br>model |
| Model resolution (Å) | 2.44 | 2.33 | 2.44 | 2.75 |
| FSC threshold | 0.5 | 0.5 | 0.5 | 0.5 |
| Map sharpening *B* factor (Å<sup>2</sup>) | -80.1 | -74.8 | -79.9 | -105.4 |
| Model composition |  |  |  |  |
| Non-hydrogen atoms | 3466 | 3516 | 3511 | 3472 |
| Protein residues | 399 | 399 | 399 | 399 |
| Ligands |  |  |  |  |
| HEM | 2 | 2 | 2 | 2 |
| FAD | 1 | 1 | 1 | 1 |
| NDP | - | 1 | - | 1 |
| NAI | - | - | 1 | - |
| Waters | 63 | 65 | 64 | 27 |
| *B* factors (Å<sup>2</sup>) |  |  |  |  |
| Protein | 34.55 | 41.92 | 37.17 | 48.66 |
| Ligand | 22.74 | 34.50 | 28.78 | 44.90 |
| Water | 31.09 | 40.97 | 35.05 | 39.25 |
| R.m.s. deviations |  |  |  |  |
| Bond lengths (Å) | 0.004 | 0.007 | 0.003 | 0.007 |
| Bond angles (°) | 0.940 | 0.870 | 0.578 | 0.872 |
| Validation |  |  |  |  |
| MolProbity score | 1.76 | 1.27 | 1.26 | 1.56 |
| Clashscore | 4.72 | 5.11 | 5.55 | 7.45 |
| Poor rotamers (%) | 0.86 | 0.86 | 0.58 | 1.16 |
| Ramachandran plot |  |  |  |  |
| Favored (%) | 98.24 | 98.24 | 98.74 | 97.48 |
| Allowed (%) | 1.76 | 1.76 | 1.26 | 2.56 |
| Disallowed (%) | 0 | 0 | 0 | 0 |
| Q-score | 0.82 | 0.83 | 0.82 | 0.77 |

### The architecture of SpNOX

The structure of SpNOX corresponds to the canonical NOX structure, with an N-terminal TM domain that coordinates two hemes, and a C-terminal domain bound to FAD (Fig. 1c-e). Despite sharing an overall sequence identity between 17 and 22% with human and cyanobacterial NOXs (Supplementary Fig. 8a), the SpNOX structure strongly resembles the structures of eukaryotic NOXs and csNOX5 TM and DH domains (Supplementary Fig. 9).

The SpNOX TMD encompasses six transmembrane helices (TM1-TM6) with an overall pyramidal shape, triangular on the inner membrane side and narrower towards the extracellular space. This folding strongly resembles the ferric reductase domain of eukaryotic NOXs, with TMs 2-5 adopting an hourglass-shaped conformation that binds two b-type hemes orthogonal to the membrane plane, one located closer to the cytosolic side (inner heme) and the other closer to the outer side (outer heme). Both are hexacoordinated via two pairs of histidines (His69 and His129, inner heme; His83 and His142, outer heme) belonging to TM3 and TM5 (Fig. 2a,b and Supplementary Fig. 8b). The iron-to-iron distance is 21.47 Å, while the edge-to-edge distance is 9.84 Å.

**Fig. 2.**
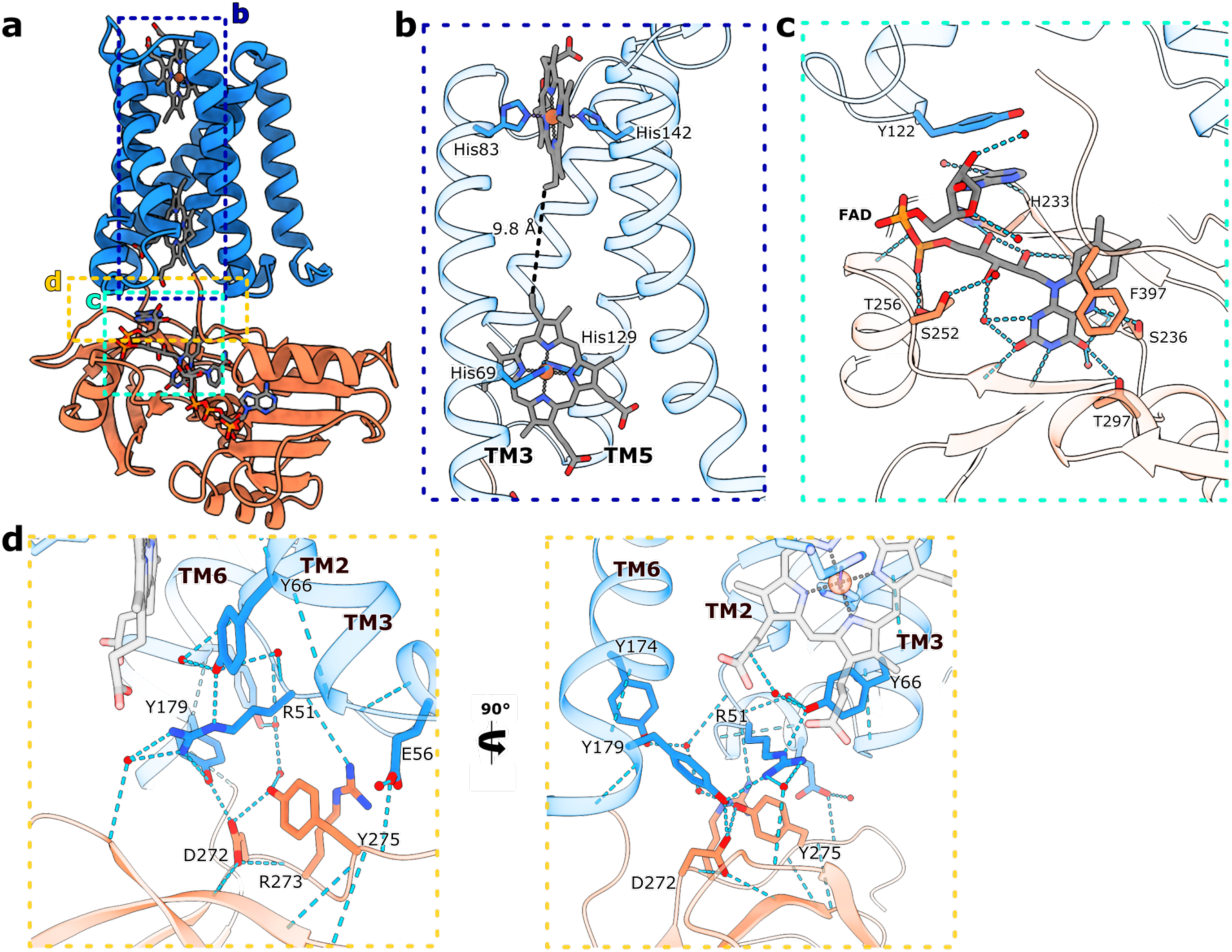
Heme-and FAD-binding sites and interactions between the TM and DH domains. **a,** Cartoon representation of the structure of NADPH-bound spNOX. Areas of interest are highlighted with colored dashed boxes. **b**, The two b-type hemes are coordinated via two histidine pairs His83/His142 (outer heme) and His69/His129 (inner heme) of the transmembrane helices TM3 and TM5. The edge-to-edge distance is indicated with a dashed line. TM1 and TM2 are omitted for clarity. **c,** Detailed view of the FAD-binding site. Side chains and FAD are shown as sticks. **d**, Closer look into the interface interactions between the TM and the DH domains. In c and d, water molecules are shown as red spheres, and atoms within H-bond distance are marked with cyan dashed lines.

The C-terminal DH domain is connected to TM6 by a short linker and shows the canonical ferredoxin-NADP^+^ reductase (FNR) folding: an FAD-binding lobe consisting of six anti-parallel β-strands forming a β-barrel flanked by an α-helix; and an NAD(P)H-binding lobe with a Rossman-like topology formed by a parallel β-sheet sandwiched between α-helices (Fig. 2a and Supplementary Fig. 10a). When compared to eukaryotic NOXs and csNOX5 (Supplementary Fig. 8c and Supplementary Fig. 9), the DH domain of SpNOX shows up to 27% sequence identity with a simplified overall architecture. The main structural differences accumulate at the NADPH-binding lobe (Supplementary Fig. 9), where SpNOX shows an unstructured long linker connecting the parallel β-strands. In eukaryotic NOXs, insertions and rearrangements at this site serve important regulatory functions (e.g. the calmodulin-binding region of NOX5). A clear density for FAD was well resolved in all maps (Supplementary Fig. 7) inside a positively charged pocket at the interface of the FAD-binding lobe and the TM domain (Supplementary Fig. 10b). The DH domain residues His233, Ser236, Lys250, Ser252, Thr256, Thr297 interact with FAD via H-bonds, whereas Phe397 interacts via π–π stacking with the isoalloxazine ring (Fig. 2c and Supplementary Fig. 10c). Residues His233 and Ser236 belong to the strictly conserved motif ‘HPF(S/T)’ (Supplementary Fig. 8c). The only direct interaction between FAD and the TM domain is through π–π stacking between the adenine and Tyr122 (Fig. 2c and Supplementary Fig. 10d), which is strictly conserved in bacterial NOX-like proteins (Supplementary Fig. 11). The DH domain is docked through direct polar interactions of residues at the FAD-binding lobe, mainly with residues located at the B-loop and TM6 of the TM domain (Fig. 2d). The adopted geometry of FAD is the same as in NOX2, which achieves this interaction by the Tyr122-equivalent residue Phe202 (Supplementary Fig. 10e). Interestingly, this FAD conformation is also present in human DUOX1, in spite of the fact that different interactions (mainly salt bridges between the AMP phosphate with Arg1214 and Arg1131 of the TM domain) are responsible for forming the interaction with FAD (Supplementary Fig. 10f). This conservation of FAD structure in NOXs, in spite of a change in the FAD-interacting amino acids, may point to an important role of this compact flavin conformation, in contrast to the commonly seen extended conformation in other flavoproteins (e.g. PDB:1B2R, 6HCY, 6LUM)[12–14].

### The NAD(P)H-binding site and the electron transfer pathway of SpNOX

NADPH-bound (2.2 Å resolution) and anaerobic NADH-bound (2.4 Å resolution) SpNOX maps showed well-resolved densities for the substrate (Fig. 3 and Supplementary Fig. 7). Aside from the bound substrate, all three structures appear largely the same with only minor side chain rearrangements (Supplementary Fig. 12a-c, e-g, i-k). The nicotinamide moiety of NADPH and NADH is accommodated via H-bonds and hydrophobic interactions inside a cavity generated by Phe397 at the C-terminal tail, and the strictly conserved ‘XGXGX’ and ‘CG(S/P)’ motifs located at the loops connecting α2 and β8, and β10 and α3, respectively (Fig. 3a and Supplementary Fig. 8c, 12d,h,l and 13a). A Cys370Ala mutant showed an increase in K_m_, confirming the relevance of the ‘CG(S/P)’ motif for substrate binding (Supplementary Fig. 14a).

**Fig. 3.**
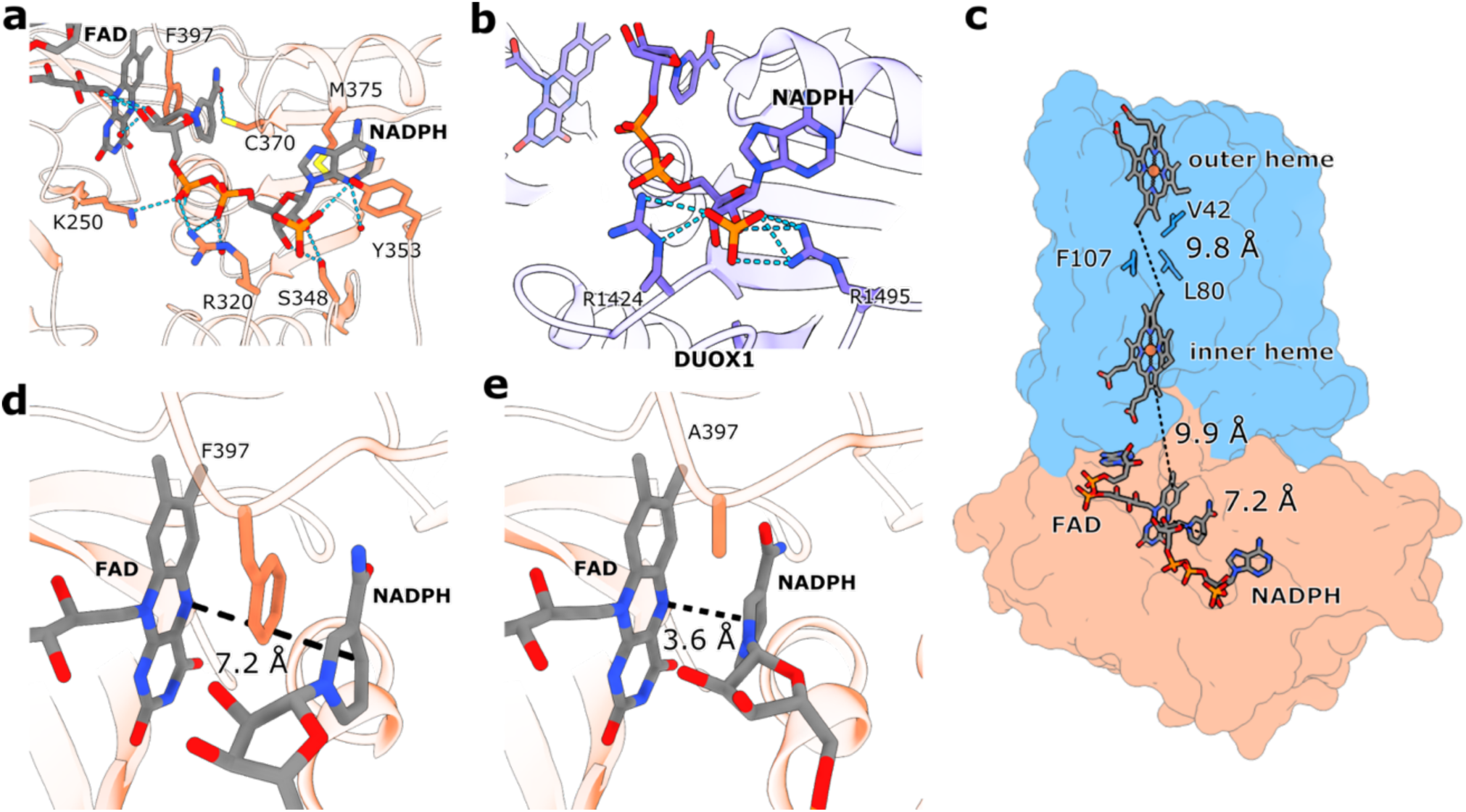
The NAD(P)H-binding site and the SpNOX electron transfer pathway. **a**, Detailed view of the NADPH-binding site at the DHD of SpNOX. Atoms within H-bond distance are marked with cyan dashed lines. FAD (grey) and amino acid side chains (coral) are shown as sticks. **b,** The lack of specificity towards NADPH in SpNOX can be explained by the absence of ionic interactions with the 2’-phosphate, contrary to eukaryotic NOXs including human DUOX1, in which Arg1412 and Arg1495 interact with the 2’-phosphate. **c,** The proposed electron transfer path within SpNOX. Hemes, FAD, NADPH and the inter-heme hydrophobic residues are shown as sticks on the surface of SpNOX. Distances between the redox cofactors are represented as dashed black lines, and were measured between the nicotinamide and the isoalloxazine ring (7.2 Å), the isoalloxazine ring and the inner heme lower edge (9.9 Å), and between the edges of the inner and outer hemes (9.8 Å). **d**, Phe397 sits between the isoalloxazine ring of FAD and the nicotinamide ring of NAD(P)H, impeding hydride transfer. **e,** The nicotinamide ring of NADPH moves closer to the isoalloxazine group of FAD in the Phe397Ala SpNOX mutant.

Overall, the AMP-binding site of both substrates shows the highest number of interactions with SpNOX. Previous structural studies of NADPH-specific FNR superfamily members like DUOX1 [9], *Anabaena* PCC 7119 FNR [15] and human neuronal nitric oxide synthase (nNOS) [16], have shown that the AMP-binding region is critical for coenzyme specificity due to the presence of at least one Arg residue that establishes charge-to-charge interactions with the 2’-phosphate group of NADPH. In SpNOX, this group is coordinated via H-bonds with Ser348 and Tyr353 (Fig. 3a), which is remarkably different to the NADPH-binding mode of eukaryotic NOXs and other members of the NADPH-specific FNR superfamily. In human DUOX1, two arginine residues, one analogous to Arg320 of SpNOX (Arg1424), and another analogous to Tyr353 (Arg1495), both coordinate the 2’-phosphate via salt bridges (Fig. 3b). In human NOX2, Arg513 also occupies the position of Tyr353 (Supplementary Fig. 8c). In *Anabaena* FNR, a Tyr residue analogous to Tyr353 interacts with the adenine moiety, but as in eukaryotic NOXs, the 2’-phosphate is also coordinated by salt bridges with two arginine residues (Supplementary Fig. 13b). Therefore, NADPH-specificity results, at least partially, from ionic interactions that are absent in SpNOX, which would explain the lack of substrate specificity. Supporting this, a Tyr353Arg mutant increased the specificity of SpNOX for NADPH by 20-fold (Supplementary Fig. 14b,c). Additionally, the 5’-phosphate interacts via a salt bridge with Arg320, the adenine moiety is sandwiched between Met375 side-chain and the aromatic ring of Tyr353, and the 3’-OH of the ribose performs H-bonding with Ser318 and Ser348. The nicotinamide-bound ribose is hydrated by several resolved water molecules but does not show any direct interaction with SpNOX, whereas the adjacent phosphate group is mainly stabilized by a salt bridge with Lys250 (Supplementary Fig. 13c).

The apparent electron transfer pathway of SpNOX corresponds to the previously described electron pathways of NOX proteins: NAD(P)H à FADà Inner Hemeà Outer Hemeà O_2_ [7–11]. The edge-to-edge distance from FAD to the inner-heme, and inner-heme to outer-heme, is 12.6 and 9.8 Å, respectively (Fig. 3c). The space between the two hemes is partially occupied by a cluster of hydrophobic amino acids which includes the aromatic residue Phe107. This residue occupies a similar position to a conserved aromatic residue present in other NOXs (Phe215 in human NOX2, Phe1097 in human DUOX1 and Trp378 in csNOX5). Mutational analysis in mouse DUOX1 and NOX2 [8, 11] showed that this amino acid could be the preferred route for electron transfer between the hemes, although the distance between both hemes is in the 14 Å range for efficient electron tunneling in all NOX structures [17]. Here, we analyzed the activity of a Phe107Leu SpNOX mutant (Supplementary Fig. 14d). Contrary to the previous observations in eukaryotic NOXs, we did not detect any significant reduction of SpNOX activity. This result indicates that in SpNOX, Phe107 is not required for efficient electron transfer and suggests that the loss of activity in other NOXs could be the consequence of a less optimal hydrophobic environment or structural rearrangements.

Our substrate-bound structures represent an inactive state of the enzyme, as deduced by a 7.2 Å core-to-core distance between the nicotinamide C4 and the isoalloxazine N5, too long for efficient hydride transfer (Fig. 3c,d), which should require distances consistent with simultaneous bond breakage and formation in the transition state [18–20]. Furthermore, Phe397 is stacked between the nicotinamide and the isoalloxazine rings (Fig. 3d), suggesting that during turnover a conformational change involving, at least, the displacement of Phe397 lateral chain is required for hydride transfer. Remarkably, an analogous C-terminal aromatic residue is conserved within other members of the FNR superfamily including plant-type FNRs, nitric oxide synthases, cytochrome P450 reductases and phthalate dioxygenase reductases [16, 21–23]. Extensive structural and biochemical studies have suggested that the displacement of this residue is highly thermodynamically unfavored and may act as the rate-limiting step for flavin reduction [21, 24–26]. This residue may also contribute to the regulation of NAD(P)H-binding affinity and specificity, and to the stabilization of the FAD semiquinone state [24]. Mutation of the C-terminal Tyr of plant-like FNRs to a non-aromatic residue like Ala or Ser substantially increases the affinity for NAD(P)^+^/NAD(P)H and induces an enzyme state with productive flavin–nicotinamide interaction [20]. However, in other FNR superfamily members like human nNOS, this kind of mutation does not lead to a large change in the binding affinity for NAD(P)H, though they increase the k_cat_ for NADH [22]. Here, we analyzed the steady state kinetics of a Phe397Ala SpNOX mutant (Supplementary Fig. 14e). Similarly to nNOS, we did not observe a large effect on the K_m_, which suggests that Phe397 does not play a major role in the control of NADPH-binding affinity in SpNOX. We observed a moderate increase in k_cat_, which could be explained by the elimination of the Phe397 displacement step. These data are supported by further analysis of a cryo-EM structure of the Phe397Ala mutant bound to NADPH under turnover conditions at 2.64 Å resolution (Table 1, Fig.3e and Supplementary Fig. 5). A density can be observed for the bound substrate but, compared to the other substrate-bound structures described here, the nicotinamide is closer to the isoalloxazine ring (Fig. 3e, Supplementary Fig. 12i-k). The absence of the large side chain of Phe397 allows NADPH to take up a productive conformation in SpNOX, as has been previously seen for pea FNR Tyr308 mutants [20]. In this conformation, the nicotinamide ring does not lie parallel to the isoalloxazine ring, but at a ∼26° angle (Fig. 3e).

A search for potential oxygen reduction centers close to the outer heme binding pocket revealed a strikingly high exposure of the outer heme to the extracellular space, which in other NOXs is more buried within the structure by the TM helices (csNOX5) [7], or by long extracellular loops (NOX2) or domains (DUOX1) acting as caps (Fig. 4a) [8, 11]. In SpNOX, the extracellular loops (A-, C-, and E-loops) fold away from the heme cavity, directly exposing the outer heme to the solvent (Fig. 4b,c) and creating a unique environment when compared to other NOXs. In fact, although all putative oxygen reaction centers previously described for NOXs showed a similar conformation involving two histidine residues, an arginine and a propionate group of the outer heme [8–11], we did not find any similar O_2_-binding site in SpNOX. However, a closer examination of the outer heme-binding pocket revealed a small cavity below the C-loop occupied by an ordered water molecule. This water molecule, which forms H-bonds with Asn84, Asn101 and Tyr105, could be occupying an O_2_-binding site within efficient electron transfer distance to the outer heme (Fig. 4d). A potential path for O_2_/O_2_^•−^ entry and exit at this site was found using Hollow [27], taking the modelled water molecule as the starting point (Supplementary Fig. 13d). However, Asn84Ala and Tyr105Phe mutants did not display a significant change in cytochrome c reduction or NADPH oxidation activity compared to WT (Supplementary Fig. 14f-h), suggesting this site may not be an actual oxygen-reaction center. We also identified another putative O_2_-binding site composed of Ser83 and heme-coordinating His93. Both amino acids are highly exposed to the solvent and coordinate a water molecule that could be occupying the O_2_-binding position (Fig. 4e).

**Fig. 4.**
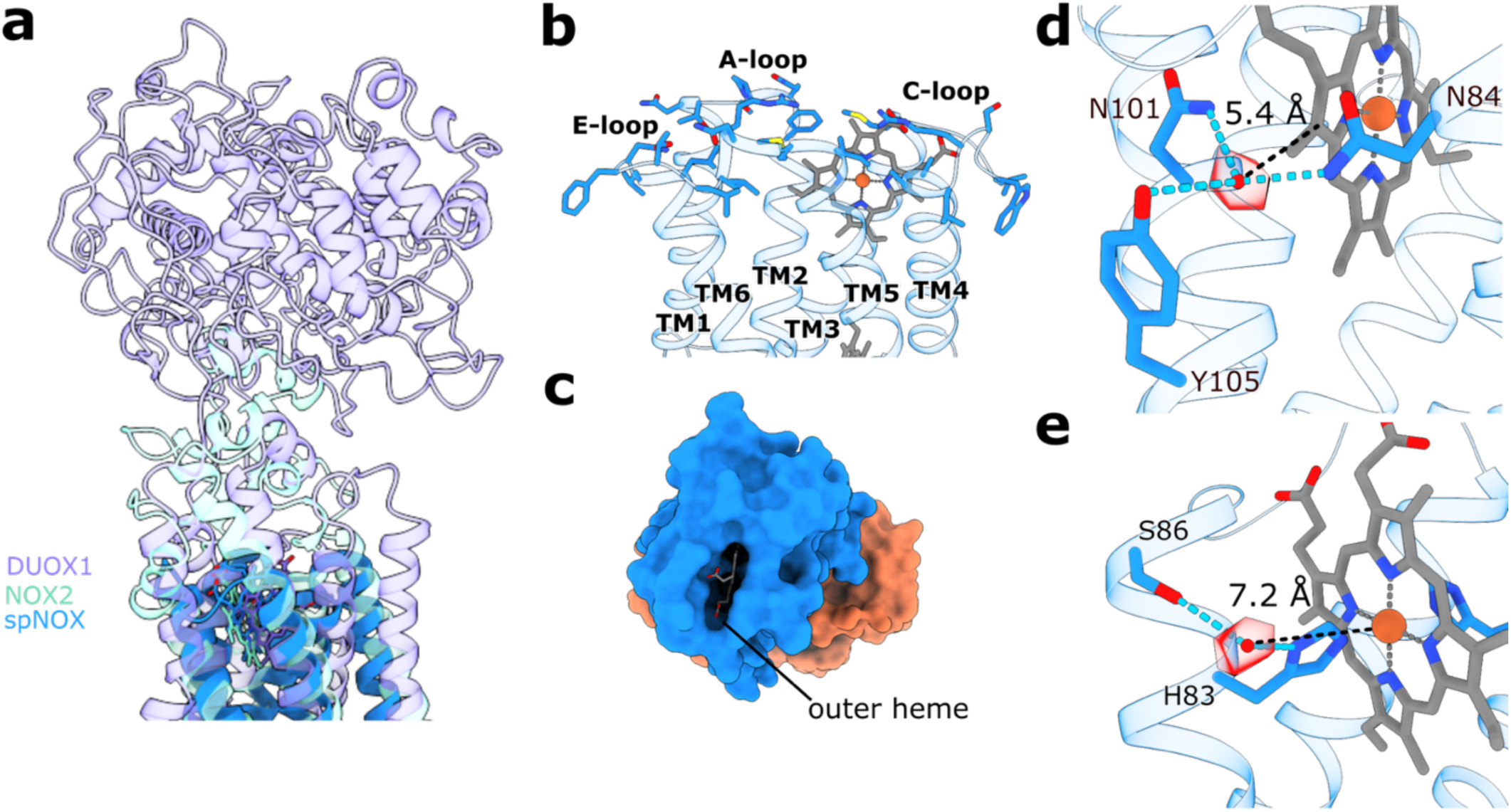
SpNOX displays a highly solvent-exposed outer heme and two putative oxygen-binding sites. **a,** SpNOX (cartoon representation, blue) presents an outer heme more solvent-exposed than eukaryotic NOXs like NOX2 (cyan) or DUOX1 (purple) due to a lack of extracellular domains or large loops. **b,** Side view of the extracellular loops of SpNOX. **c,** Top view of SpNOX surface showing the solvent-exposed outer heme. **d & e,** Potential O_2_-reduction centers. The distance between the coordinated water molecules (red sphere and density) and the outer heme is shown as a black dashed line. Atoms within H-bond distance are marked with cyan dashed lines.

## Discussion

Phylogenetic analyses of the FRD-containing superfamily of proteins indicate that the NOX family emerged early in evolution by the fusion of a bacterial FRD-containing protein with an FNR protein [5]. This hypothesis has been supported by the recent discovery of NOX-like homologues in bacteria, and the biochemical characterization of a member of this new group, SpNOX [4, 6]. The cryo-EM structures of this protein presented here provide further relevant details about its catalytic mechanism and add new insights into the electron transfer pathway of NADPH oxidases.

*S. pneumoniae* is a Gram-positive, aerotolerant anaerobic bacterium that is part of the human nasopharyngeal microbiota but can cause serious infections [28]. The mechanisms by which it deals with oxygen and ROS have been extensively studied [29–32]. Remarkably, *S. pneumoniae* is able to produce large amounts of H_2_O_2_ that appear to promote infection by damaging epithelial cells and other microorganisms of the respiratory tract [33, 34]. Nevertheless, contrary to eukaryotic cells, which tightly regulate ROS production by NOXs, *S. pneumoniae* produces H_2_O_2_ as a metabolic byproduct of enzymes like the pyruvate oxidase SpxB [30]. Whether bacteria use SpNOX-like proteins for dedicated ROS production *in vivo* is unresolved. An SpNOX-knockout strain of *S. pneumoniae* was reported to display no discernible phenotype in pure culture [4].

An important characteristic of SpNOX is its constitutive activity *in vitro,* a feature only shared by mammalian NOX4 homologues. The exact structural basis for this feature remains unknown, but published cryo-EM structures of eukaryotic NOX show a highly flexible DH domain whose conformational state is believed to respond to regulatory signals (e.g. calcium in DUOX1 or cytosolic factors in NOX2) [9, 11]. The structures we obtained suggest that, in SpNOX, the DH domain forms very stable contacts with the TM domain even in the absence of substrate or native lipids, as deduced by the high-resolution achieved for both DH and TM domains in a consensus refinement. Therefore, the stability of the interaction of the DH and the TM domains could be a major mechanism to achieve constitutive activity in NOX proteins. Our substrate-bound structures both reveal an unproductive conformation for hydride transfer, providing important detail of the catalytic mechanism of SpNOX. Like for other members of the FNR superfamily, a transient displacement of a C-terminal aromatic residue is necessary to perform hydride transfer [24, 25]. This is supported by an increase of the turnover rate in an SpNOX Phe397Ala mutant for which our structure shows a productive conformation of the nicotinamide (Supplementary Fig. 12 and Fig. 3d,e). Interestingly, a similar catalytic role has been proposed for a strictly conserved Phe residue at the C-terminus of eukaryotic NOXs [21]. However, none of the published structures could show this residue interacting with FAD. In csNOX5, the hyperstabilizing mutation used for crystallization of the DH domain seems to prevent Phe693 from folding natively [7]. In inactive NOX2 complexes, the C-terminal residues could not be resolved due to flexibility [10, 11]. In the high-calcium state structure of human DUOX1, Phe1551 does not interact with FAD and adopts a different conformation far away from the nicotinamide ring. Nevertheless, the distance between the nicotinamide and the isoalloxazine rings is longer than 10 Å, indicating that DUOX1 activation could involve further conformational changes [9]. Therefore, with the structural information currently available, it cannot be excluded that the C-terminal Phe of eukaryotic NOXs plays a similar role to SpNOX Phe397.

Despite the large evolutionary distance between eukaryotic NOXs and SpNOX-like bacterial NOXs, our structural and biochemical analyses of SpNOX reveal a highly similar domain organization and fold. This indicates that preserved structural motifs like the hydrophobic cleft between the inner and the outer heme could be essential for efficient function of the complex. SpNOX also displays relevant specific features not present in eukaryotic NOXs. For example, it lacks the canonical amino acids that configure the potential oxygen reduction center in the solved structures of eukaryotic and cyanobacterial NOXs. Structural analysis revealed two putative oxygen reduction centers (Fig. 4d,e) with an unusual configuration when compared to other reduction centers previously described in flavoenzymes with oxidase activity [35]. These amino acids are not strictly conserved in other putative bacterial NOXs (Supplementary Fig. 11), but a similar amino acid configuration to the putative center formed by Asn84, Asn101 and Tyr105 could be found in several SpNOX-like proteins by analysis of the Alpha-Fold2 (AF2) [36] predicted structures (Supplementary Fig. 15). However, mutational analysis of these residues did not show a reduction of the activity (Supplementary Fig. 14f-h), indicating that NADPH-oxidase activity of SpNOX could be independent of this site. Therefore, we cannot exclude that oxygen is not the physiological electron acceptor of bacterial SpNOX homologues. The highly exposed outer heme of SpNOX is a feature shared with the protein-methionine-sulfoxide reductase heme-binding subunit MsrQ [37]. MsrQ is considered an ancestor of the FRD superfamily and is part of the MsrPQ system of Gram-negative bacteria [5], transferring electrons from the quinone pool to the periplasmic subunit MsrP, which finally reduces methionine sulfoxide residues of periplasmic proteins . Although the SpNOX TM domain shares only 22% identity with *E. coli* MsrQ, a comparison between the *E. coli* MsrQ AF2 model and substrate-free SpNOX revealed a similar structure (Supplementary Fig. 16). Remarkably, the extracellular region of MsrQ, which is the interaction site of the electron acceptor MsrP, is formed by short loops and is solvent-exposed at the heme-binding cleft. The fact that SpNOX shows an even more solvent-exposed extracellular region (Fig. 4b,c and Supplementary Fig. 16c-e) could indicate that a potential protein acceptor exists for this protein. This would also be supported by the fact that SpNOX can directly reduce cytochrome c and SOD efficiently, as deduced by our (an)aerobic activity assays (Supplementary Fig. 1e,f), and by previous assays under an argon atmosphere [4]. Another relevant characteristic of SpNOX is its lack of specificity towards NADPH or NADH. Further functional assays are needed to identify the physiological role of this protein and other bacterial NOX homologues.

A major achievement of our study is the determination of high-resolution cryo-EM structures of SpNOX despite its relatively small size (46 kDa). The fact that we were able to obtain high-resolution reconstructions of such a small membrane protein without the use of a fiducial (e.g. a protein-specific nanobody) illustrates the impact of ongoing developments in microscope, detector and software capabilities, that continue to push the boundaries of what is possible by single-particle cryo-EM. Important factors in our result are likely to include the production of a highly stable and homogeneous sample (Supplementary Fig. 1a,b); extensive sample optimization for optimal particle distribution in thin ice (Supplementary Figs. 2-5); the use of a Krios G4 microscope with cold field emission gun for data collection; a highly stable and low-aberration energy filter [38]; high-DQE Falcon 4 detector [39]; the use of holey gold grids to reduce beam-induced motion [40]; and a partially rigid structure that allows particle alignment.

These high-resolution structures, in substrate-free and stably reduced forms, and under turnover conditions, reveal important new detail about the catalytic activity and the electron-transfer pathway of this enzyme, shedding light on bacterial NOX homologues and similarities and differences to eukaryotic NOXs. A complete structural understanding of this emerging model enzyme informs functional studies and drug-discovery experiments involving NOX proteins.

## Methods

### Protein expression and purification

An *E. coli* codon-optimized sequence encoding the *Streptococcus pneumoniae* R6 strain NOX (2-400, Q8CZ28_STRR6), with an N-terminal 6xHis-tag followed by a PreScission cleavage site, was purchased from GenScript and subcloned between the NcoI and EcoRI sites of the pET-28a plasmid. For the generation of mutants, primers with overlapping sequences (Supplementary Tables 1 and 2) were designed to generate point mutations with the NEBuilder HiFi DNA Assembly mix (New England Biolabs). Wild-type protein expression was performed in OverExpress C41 (DE3) *E. coli* cells. Cultures grown in Terrific broth (TB) at 37°C were induced in the exponential phase (OD600 = 1.0) by the addition of 0.2 mM Isopropyl ß-D-1-thiogalactopyranoside (IPTG). The cultures were supplemented with 0.5 mM δ-aminolevulinic acid (TCI) to maximize heme incorporation. After 4 hours at 37°C and 140 rpm, the cells were harvested and flash frozen in liquid N_2_ for later use. For protein purification, the frozen cell pellets were thawed on ice and resuspended in lysis buffer containing 50 mM Tris-HCl pH 7.0 and 300 mM NaCl supplemented with bovine DNase II, 1 mM PMSF and 5 µg/mL aprotinin, leupeptin and pepstatin (10 mL per 1 g cell pellet) at room temperature for 30 minutes. Resuspended cells were lysed on ice using an ultrasonic homogenizer (Bandelin) with 3 cycles at 40% amplitude with one second on and one second off for a total of 6 minutes. All subsequent steps were performed at 4°C. The cell debris was pelleted by centrifugation at 12,108 x g for 30 minutes. The membranes were isolated from the supernatant by ultracentrifugation at 185,511 x g for 1 hour. Then, membrane pellets were resuspended in a Dounce homogenizer in lysis buffer. Once resuspended the sample was supplemented with 0.4% LMNG and gently stirred overnight for solubilization. Non-solubilized material was removed the next day by ultracentrifugation at 185,511 x g for 30 minutes. The supernatant was mixed with Ni-NTA resin equilibrated in lysis buffer and incubated for 2 hours while gently rocking. The incubated resin was then loaded onto a gravity column and washed with 10 column volumes of lysis buffer supplemented with 5 mM imidazole and 0.002% LMNG. The protein was eluted with 300 mM imidazole and immediately diluted 1:1 with imidazole free lysis buffer supplemented with 0.002% LMNG to avoid aggregation. PD-10 columns (GE Healthcare) were used for buffer exchange to 50 mM Tris-HCl pH 7.5, 150 mM NaCl and 0.002% LMNG. His-PreScission protease, expressed and purified in-house [41] was added to a final concentration of 0.05 mg/mL and incubated overnight. The sample was centrifuged at 12,108 x g for 30 minutes to pellet aggregates, supplemented with 20 mM imidazole and loaded onto a Ni-NTA gravity column equilibrated in 50 mM Tris-HCl pH 7.5, 150 mM NaCl, 20 mM imidazole and 0.002% LMNG. The flow-through volume containing the untagged protein was concentrated to 10 mg/mL using Vivacon500 filters (50 kDa, Sartorius) for further purification using a Superdex 200 Increase 5/150 GL column equilibrated in 50 mM Tris-HCl pH 7.0, 250 mM NaCl and 0.002% LMNG. Fractions containing the monodisperse protein were pooled, concentrated for cryo-EM sample preparation or flash frozen for future use.

### Cryo-EM sample preparation and data collection

All cryo-EM grids were prepared from freshly purified SpNOX concentrated to 6 mg/mL for substrate absent samples and 7.5 mg/mL for substrate present sample and supplemented with 1 mM FAD at least 30 minutes before grid freezing.

For the substrate-free structure, 3 µL SpNOX was directly applied to freshly glow-discharged 1.2/1.3 400 mesh UltrAuFoil grids. The grids were blotted using Whatman 595 blotting paper (Sigma-Aldrich) for 4 to 6 seconds at 4°C and 100% humidity, and then plunge-frozen into liquid ethane using a Vitrobot Mark IV (Thermo Fisher Scientific). A dataset was collected using a Titan Krios G4 equipped with a cold field emission gun, Selectris X filter and Falcon 4 direct electron detector. EPU was used for automated data acquisition of 18,433 movies in EER format [42]. Movies were collected with a pixel size of 0.573 Å and a total dose of 70 e^-^/Å^2^ contained in 875 EER frames. The defocus ranged from −0.7 to −2.1 µm.

For the NADPH-bound structure, grids were prepared as for the substrate-free SpNOX except that 2.7 µL SpNOX was mixed with 0.3 µL 10 mM NADPH and immediately applied to glow-discharged 1.2/1.3 400 mesh UltrAuFoil grids inside the Vitrobot chamber. The amount of time from mixing to plunge-freezing was not longer than 15 s. A dataset was collected using the same microscope and imaging settings as for the substrate-free sample. The dataset contained 22,259 movies with 1,036 EER frames.

For the NADH-bound structure, an aliquot of aerobically purified SpNOX was put inside an anaerobic chamber (<1 ppm O_2_; Coy Laboratory Products) and mixed with 5 mM NADH from a 50 mM NADH stock. After 40 minutes, 3 µL of the sample was applied to glow-discharged 1.2/1.3 400 mesh UltrAuFoil grids, blotted for 4 to 5 seconds at 4°C under 100% humidity, and plunge-frozen in liquid ethane using a Vitrobot Mark IV (Thermo Fisher Scientific). A dataset was collected using the same microscope and imaging settings as for the aerobically frozen samples. The dataset contained 28,143 movies with 938 EER frames.

For the F397A mutant structure, grids were prepared as for the NADPH-bound structure. 2.7 µL SpNOX was mixed with 0.3 µL 50 mM NADPH and immediately applied to glow-discharged 1.2/1.3 400 mesh UltrAuFoil grids inside the Vitrobot chamber. The grids were blotted for 5-6 seconds and plunge-frozen in liquid ethane. Four datasets were collected on 2 grids using the same microscope and imaging settings as for the other datasets. Only during the session, plasmon imaging was used to efficiently select holes with the optimal ice thickness [43]. The datasets contained 21,851, 9,538, 6,332 and 7,415 movies with 924, 899, 1008 and 994 EER frames, respectively.

### Image processing

Detailed workflows for each dataset are shown in Supplementary Figs. 2-5. Processing of the substrate-free dataset started by importing all movies into RELION 4.0 [44] and the raw movies were fractioned to 1.04 e^-^/Å^2^/frame, motion corrected and dose weighted using RELION’s implementation of MotionCor2 with 5x5 patches [45]. CTF parameters were estimated with CTFFIND4.1 [46]. Micrographs were then selected based on a maximum 4.0 Å CTF resolution estimate and a defocus value between −0.5 and −2.5 µm. For particle picking different programs and models were used in parallel. After classification and removal of ‘junk’ particles, the particle sets from the different strategies were merged and duplicates were removed. Particles were picked with two trained Topaz models [47], as well as a refined standard crYOLO [48] model, from all motion corrected micrographs. The Topaz models were trained on a previous dataset which did not yield a high-resolution reconstruction. The crYOLO model was refined with ∼800 manually picked particles from selected micrographs of the substrate-free dataset. A picking threshold of 0.1 was used for crYOLO picking. Picked particles were extracted with a 256-pixel box size and downsampled to 56 pixels. Topaz particles were extracted with a figure of merit (FOM) threshold of −1.5. The particles were imported into cryoSPARC 3.2 [49] and subjected separately to several rounds of 2D-classification. Initial models were generated from 1,000,000 particles of the crYOLO particle set. The initial models were used as references for a heterogenous refinement for each particle set with 3 references. The classes representing the protein were transferred to RELION format before being merged to remove duplicate particles, resulting in a 1,160,039-particle dataset which was re-extracted in a box of 320 pixels downsampled to 72 pixels. Particle transfer between RELION and cryoSPARC was performed by the UCSF pyem program package [50]. Several rounds of non-uniform (NU) refinement and re-extraction in a larger particle box with less downsampling were performed followed by local and global CTF refinement which resulted in a 2.65 Å reconstruction using a 448-pixel box downsampled to 256 pixels. A large extraction box was necessary in order to capture all the delocalized signal of the protein particle. Several rounds of heterogenous refinement, CTF refinement and RELION-assisted particle polishing [51] resulted in a NU refinement resolved to 2.40 Å. A final local refinement with a tighter mask resulted in a 2.29 Å final reconstruction from 379,972 particles sharpened with a B-factor of −80.1 Å^2^.

Processing of the turnover conditions with NADPH present started by importing all movies into RELION 4.0 and the movies were fractioned to 1.04 e^-^/Å^2^/frame, motion corrected and dose weighted using RELION implementation of MotionCor2 using 5x5 patches. Subsequently, CTF parameters were estimated using CTFFIND4.1 and micrographs were selected based on a maximum of 3.5 Å CTF resolution estimate and a defocus value between −0.5 and −2.5 µm. The same three particle picking models and parameters were used as for the substrate-free dataset. The particles were extracted in a 256-pixel box and downsampled to 56 pixels. For the Topaz picked particles, a FOM threshold of −1.5 was used. The particles were imported into cryoSPARC and subjected to two rounds of 2D classification. One of the three particle sets was used to generate four initial models of which three were used for subsequent heterogenous refinement for each particle set after 2D classification. The SpNOX representing classes were transferred to RELION using the UCSF pyem program package. The particles were merged and duplicates removed before re-extraction in a larger box with smaller downsampling. The particles were then imported into cryoSPARC and another *ab initio* was done to classify the particles into two classes before several rounds of NU and CTF refinements with increasingly larger extraction box size and smaller downsampling. RELION-assisted particle polishing and several iterative rounds of heterogenous refinement resulted in a NU refinement reconstruction reaching 2.25 Å using 591,137 particles in a 448-pixel box downsampled to 280 pixels and a loose mask including the detergent micelle. A subsequent local refinement with a tight masking excluding the detergent micelle resulted in the final reconstruction reaching 2.20 Å nominal resolution sharpened with a B-factor of −74.8. Å^2^.

Processing of the dataset under stably reducing conditions with NADH started by importing the raw movies into RELION 4.0. The movies were fractioned to 1.04 e^-^/Å^2^/frame, motion corrected using 5x5 patches and dose weighted using RELIONs implementation of MotionCor2. The CTF parameters were estimated using CTFFIND4.1. Micrographs with a maximum of 3.5 Å CTF resolution estimate and a defocus value between −0.5 and −2.5 µm were selected. The same particle picking models as for the other datasets were used. Particles were extracted in a 256-pixel box and downsampled to 56 pixels. Topaz particles were extracted with a FOM threshold of −1.5. The particles were transferred to cryoSPARC for 2-3 rounds of 2D classification. Using one particle set, three initial models were generated and these were used for a heterogenous refinement of all three particles sets independently. The SpNOX representing classes were transferred to RELION using the UCSF pyem program package. The particle sets were merged and duplicate particles were removed. The resulting particle set was re-extracted with a larger box size and less downsampling, and then transferred to cryoSPARC. Several rounds of heterogenous, NU and CTF refinements, multiple re-extractions in a larger box with progressively less downsampling and RELION assisted particle polishing resulted in a final particle set of 697,514 particles. The final NU refinement with a loose mask resulted in a 2.42 Å reconstruction and the subsequent local refinement resulted in a 2.36 Å nominal resolution reconstruction sharpened with a B-factor of −79.9 Å^2^.

Processing of the four F397A mutant datasets started with the import of the raw movies into RELION 4.0 using separate optics groups. The raw movies were motion corrected and fractionated to 0.98, 1.03, 0.972 and 0.98 e^-^/Å^2^/frame, respectively, and dose weighted using RELION’s implementation of MotionCor2. The CTF parameters of the micrographs were estimated using CTFFind4.1. A selection of micrographs was made based on 4 Å maximum CTF resolution and defocus values between −0.5 and −2.5 µm. The same particle picking models were used as for the other datasets. Particles were extracted using a 256-pixel box downsampled to 56 pixels. The particles picked using Topaz were extracted with a FOM of −1.5. All particles from different datasets picked using the same model were merged. The particle sets were imported into cryoSPARC for 2 rounds of 2D classification. Three initial models were generated and used as refences for a round of heterogeneous refinement. The SpNOX representing classes were re-imported into RELION, merged and duplicate particles were removed. The resulting particle set was subjected to iterative rounds of NU/local refinement, CTF refinement, heterogeneous refinement, re-extraction into larger boxes with less downsampling, and RELION-assisted particle polishing. The final particle set contained 546,234 particles. The final NU refinement with a loose masking including the detergent micelle reached 2.73 Å and a subsequent local refinement using a mask excluding the detergent micelle resulted in a 2.64 Å nominal resolution reconstruction sharpened with a B-factor of −105.4 Å^2^.

### Model building

For the SpNOX apo model, the AlphaFold 2 [36] predicted structure was used as an initial model. UCSF ChimeraX [52] was used to rigidly fit the model in the density map . The program COOT [53] was used to place the cofactors and to inspect and adjust all coordinates manually. Several iterative rounds of PHENIX real-space refinement [54] and manual adjustment in COOT were performed until the stereo-chemistry was good and fit the cryo-EM density map as assessed with PHENIX, MolProbity [55] and Q-score [56].

The substrate bound models were built using the substrate-free model as an initial model except for the F397A model, there the NADPH model was used as an initial model. The respective substrate we fit into the density using COOT. Several iterations of PHENIX real-space refinement and manual adjustment were performed until the stereo-chemistry and fit in the density map were satisfactory as assessed by PHENIX, MolProbity and Q-score.

For structural comparison, models were aligned using the Matchmaker tool in UCSF Chimera X. Cavities were calculated and drawn using Hollow [27] with a 1.1-Å probe radius. Finalized models were visualized using UCSF ChimeraX. Electrostatic potential was calculated using the Adaptive Poisson-Boltzmann Solver (APBS) tool [57] and visualized in UCSF ChimeraX. Search of bacterial NOX-like proteins was performed using the BLAST similarity search tool in UniProt [58] against the UniProtKB and Swiss-Prot databases with default settings and Bacteria (Eubacteria) as taxonomy restrictor. Sequence alignments were performed with Clustal Omega [59], and visualized using the ENDscript server [60].

### Activity assay

The NAD(P)H-oxidase activity of SpNOX was measured using the cytochrome c reduction assay as previously described [61], with some modifications. The reaction was performed at room temperature in a final volume of 0.5 mL in 50 mM Tris pH 7.0, 250 mM NaCl with 100 µM bovine heart cytochrome c (Sigma). Before the assay, the solution was supplemented with 0.1 µM SpNOX and 1 µM FAD from 10 mM and 50 µM stocks, respectively. The reaction was triggered by addition of NADPH or NADH from 25 mM stocks, and the reduction was followed by measuring the absorbance at 550 nm using a spectrophotometer (Varian Cary 50). When indicated, superoxide dismutase (bovine SOD, recombinant, Sigma) was added from a ≥50,000 U/mL stock prepared in assay buffer. Anaerobic cytochrome c and SOD reduction was measured with 0.05 µM SpNOX using a Nanodrop One with cuvette holder (Thermo Fisher Scientific). Cytochrome c was added to a final concentration of 100 µM and SOD to 1000 U/mL. The oxidation of NADPH and reduction of cytochrome c were followed by measuring the absorbance at 340 and 550 nm, respectively. Aerobic and anaerobic NADPH oxidation was measured using 0.5 µM SpNOX and the reaction was initiated by the addition of 100 µM NADPH. Comparison of aerobic and anaerobic cytochrome c reduction was measured using 0.5 µM SpNOX, 100 µM NADPH and 100 µM cytochrome c. Data were analyzed and fit using OriginLab.

## Data availability

Atomic models and maps for substrate-free SpNOX were deposited in the Protein Data Bank with accession code 8QT6 and Electron Microscopy Data Bank with accession code EMD-18644; for anaerobically frozen NADH-bound SpNOX model and map are available as 8QT9 and EMD-18646; for NADPH-bound SpNOX under turnover conditions model and map are available as 8QT7 and EMD-18645; and for NADPH-bound F397A SpNOX under turnover conditions model and map are available as 8QTA and EMD-18647. Other data and materials are available from the corresponding authors upon request. Raw data have been deposited in EMPIAR under the following accession codes: substrate-free SpNOX (), NADH-bound (), NADPH-bound SpNOX () and NADPH-bound F397A NADPH-bound SpNOX (). Source data for raw images of cropped gels, as well as individual replicates of all assay results, are provided with this paper.

## Acknowledgements

We thank the Central Electron Microscopy Facility of the Max Planck Institute of Biophysics for providing cryo-EM infrastructure and technical support, and Werner Kühlbrandt and the department of Structural Biology for support and helpful discussions. We thank Rita Zimmermann for sharing her valuable expertise in anaerobic work. This work was funded by the Max Planck Society (to B.J.M.) and the Alfonso Martín Escudero postdoctoral fellowship (to P.S-A.).

## Author contributions

P.S.-A. and B.J.M conceived of the project. P.S.-A and V.R.A.D performed the biochemical and structural experiments, and analyzed the data. P.S.-A, V.R.A.D. and B.J.M. wrote the paper. B.J.M supervised the project.

## Supplementary Information for

**Supplementary Fig. 1.**
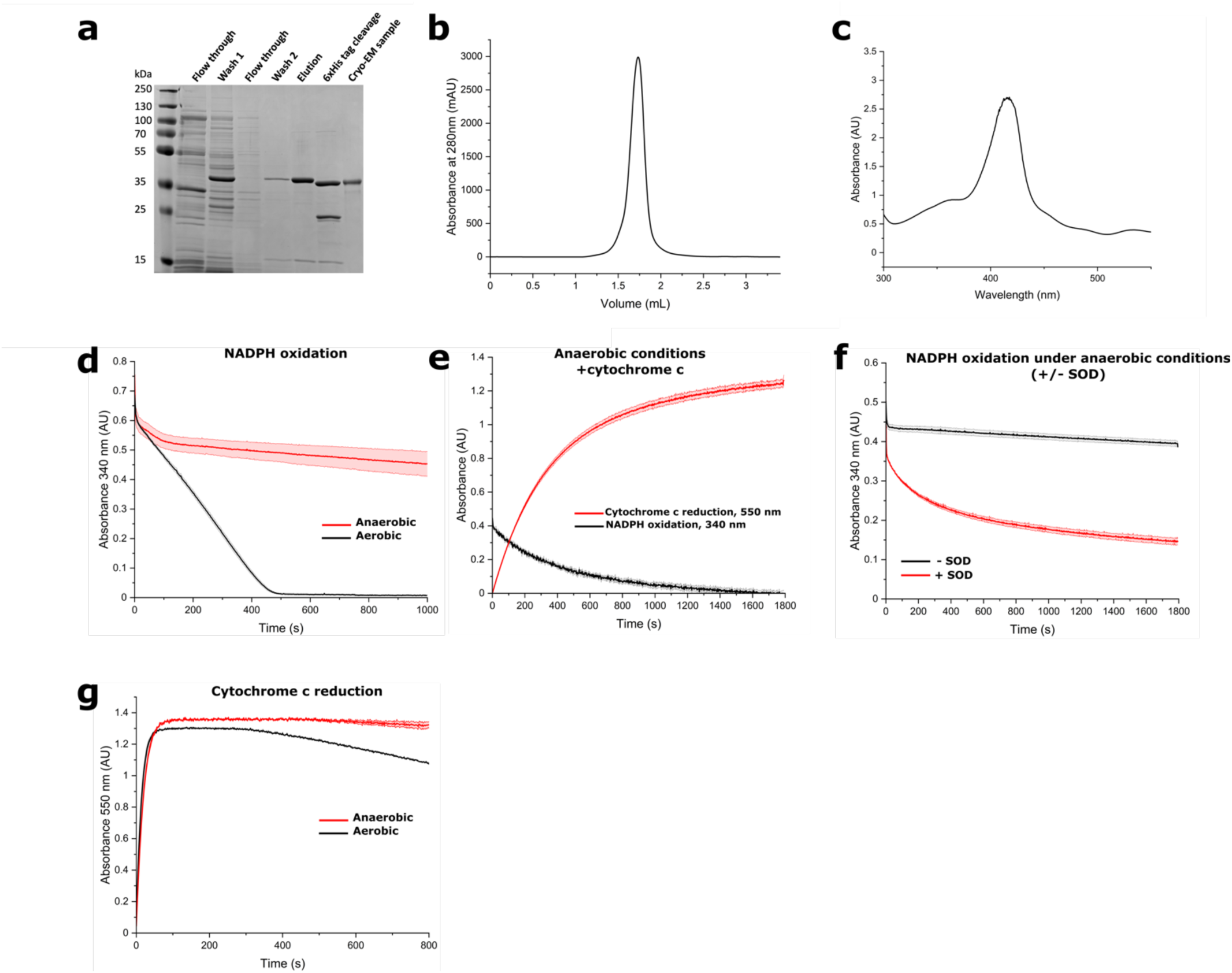
SpNOX for structural studies was highly pure and active. **a,** Analysis of the relevant fractions of SpNOX purification by Coomassie blue staining after 12% SDS-PAGE under reducing conditions. **b,** Representative size-exclusion chromatography (SEC) profile of SpNOX in LMNG micelles before cryo-EM grid freezing. **c,** UV-vis absorbance spectrum of SpNOX displaying the Soret peak at 414 nm. **D,** Directly monitoring the NADPH concentration provides a measure of cytochrome c-independent activity, which is high in the presence of oxygen (black trace), and only shows initial activity before stalling in anaerobic conditions (red trace). **e,** Anaerobic NADPH oxidation (340 nm, black trace) and cytochrome c reduction activity (550 nm, red trace) of SpNOX showing (direct) cytochrome c reduction in the absence of oxygen. **f,** Anaerobic NADPH oxidation activity of SpNOX in the presence of superoxide dismutase (SOD) showing continued NADPH oxidation in the absence of oxygen. **g,** Cytochrome c reduction activity of SpNOX shows a highly similar initial rate under aerobic (black) and anaerobic (red) conditions. Mean values of three technical replicates are plotted and SEM are indicated. Data for individual replicates are available in the source data file.

**Supplementary Fig. 2.**
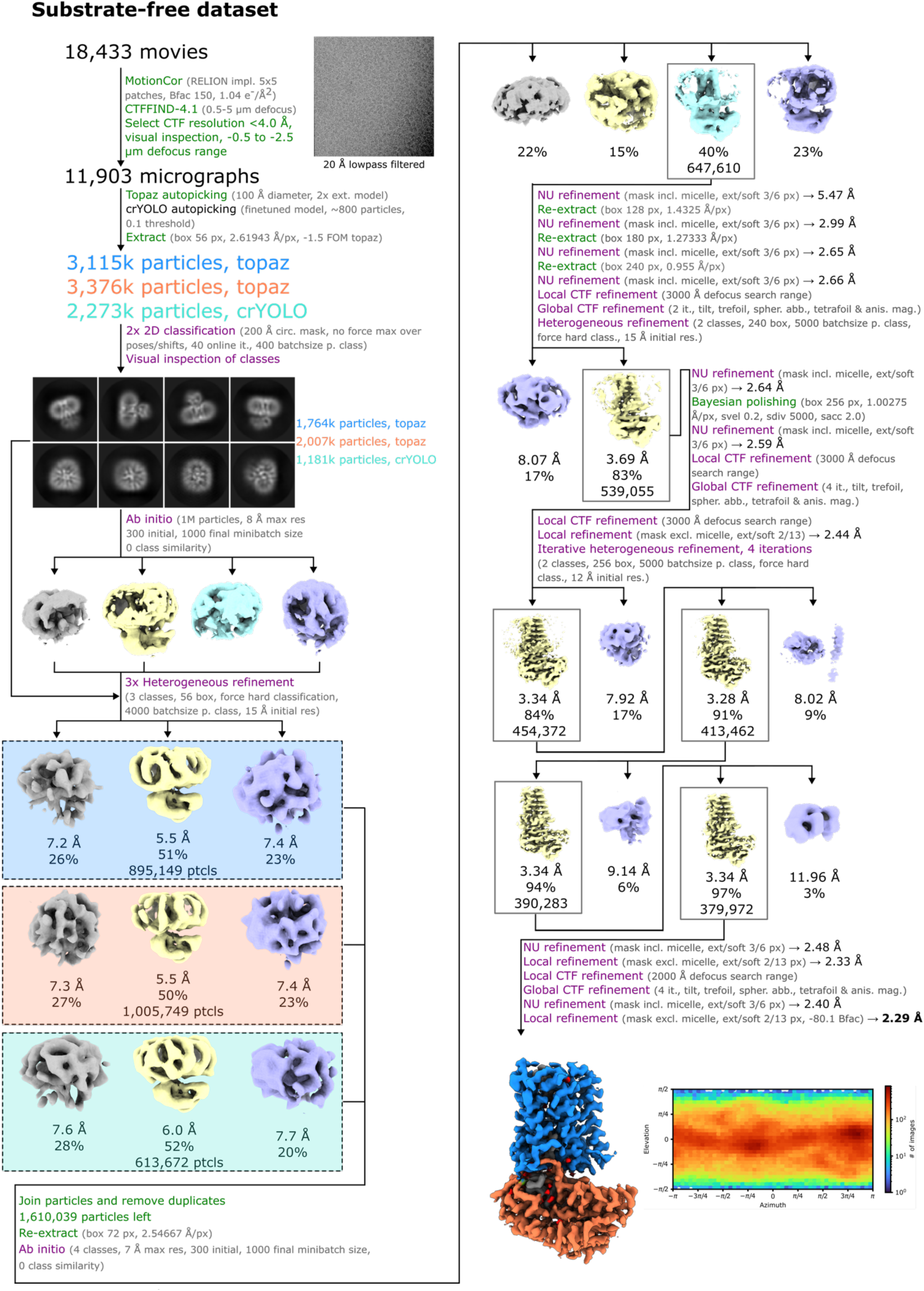
Cryo-EM processing workflow for substrate-free SpNOX reconstruction. Processes run in RELION are colored green, while processes run in cryoSPARC are colored purple. Abbreviations used: External (ext.), pixel (px), per (p.), resolution (res.), particles (ptcls), including (incl.), extension (ext), soft-edge (soft), classification (class.), iterations (it.), excluding (excl.), spherical aberration (spher. abb.), anisotropic magnification (anis. mag.) and B-factor (Bfac).

**Supplementary Fig. 3.**
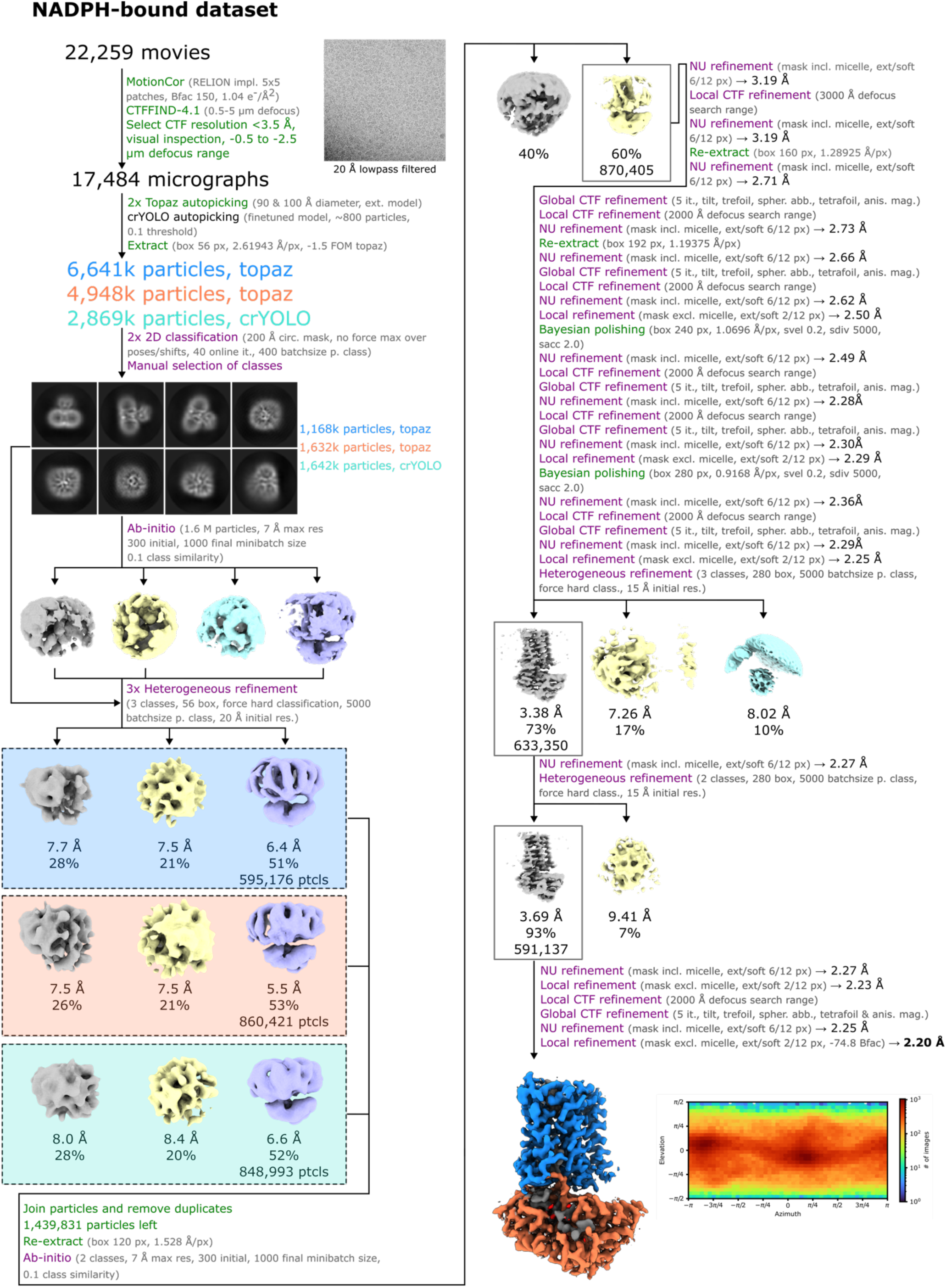
Cryo-EM processing workflow for NADPH-bound SpNOX reconstruction. Processes run in RELION are colored green, while processes run in cryoSPARC are colored purple. Abbreviations used: External (ext.), pixel (px), per (p.), resolution (res.), particles (ptcls), including (incl.), extension (ext), soft-edge (soft), classification (class.), iterations (it.), excluding (excl.), spherical aberration (spher. abb.), anisotropic magnification (anis. mag.) and B-factor (Bfac).

**Supplementary Fig. 4.**
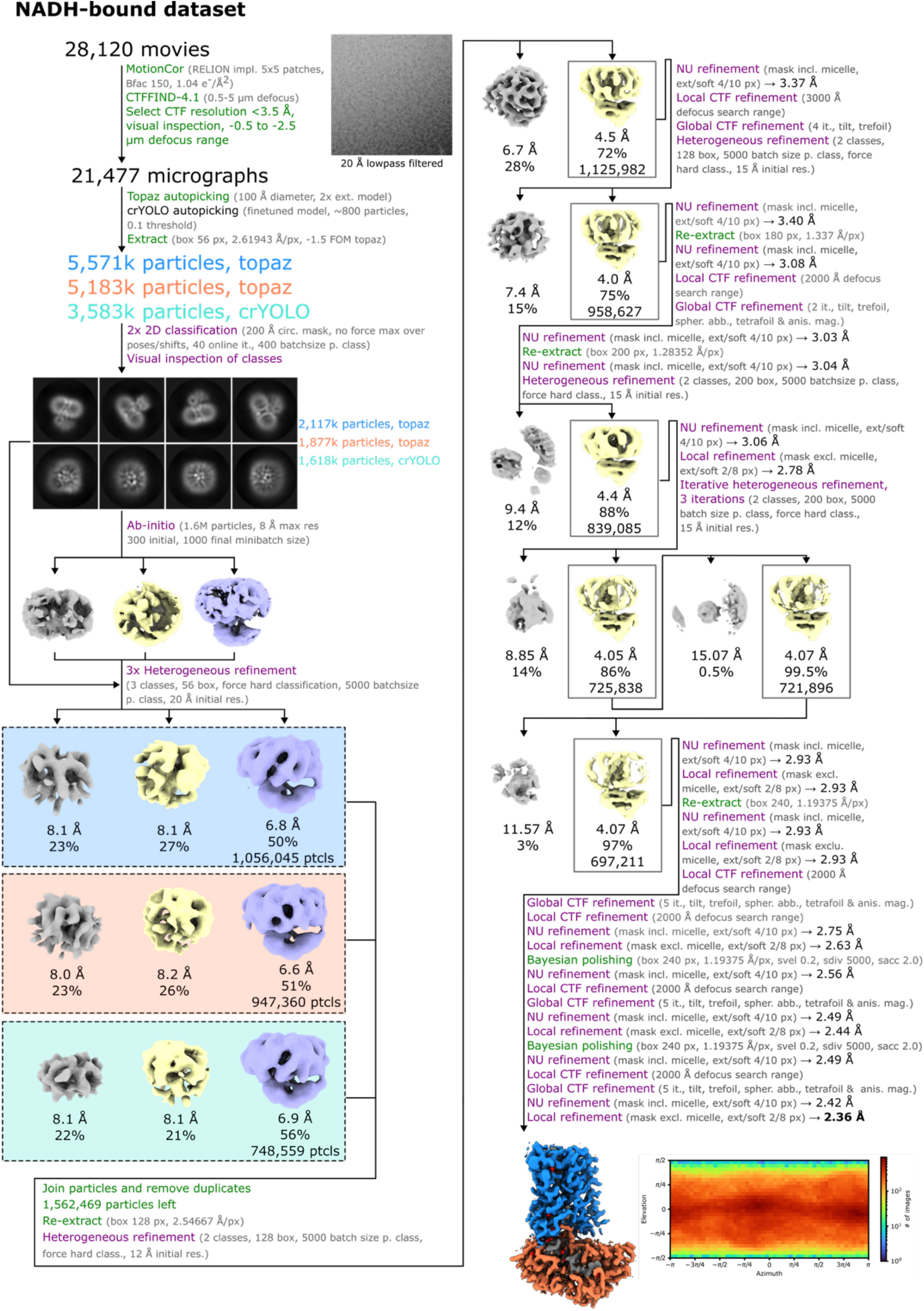
Cryo-EM processing workflow for NADH-bound SpNOX reconstruction. Processes run in RELION are colored green, while processes run in cryoSPARC are colored purple. Abbreviations used: External (ext.), pixel (px), per (p.), resolution (res), particles (ptcls), including (incl.), extension (ext), soft-edge (soft), classification (class.), iterations (it.), excluding (excl.), spherical aberration (spher. abb.), anisotropic magnification (anis. mag.) and B-factor (Bfac).

**Supplementary Fig. 5.**
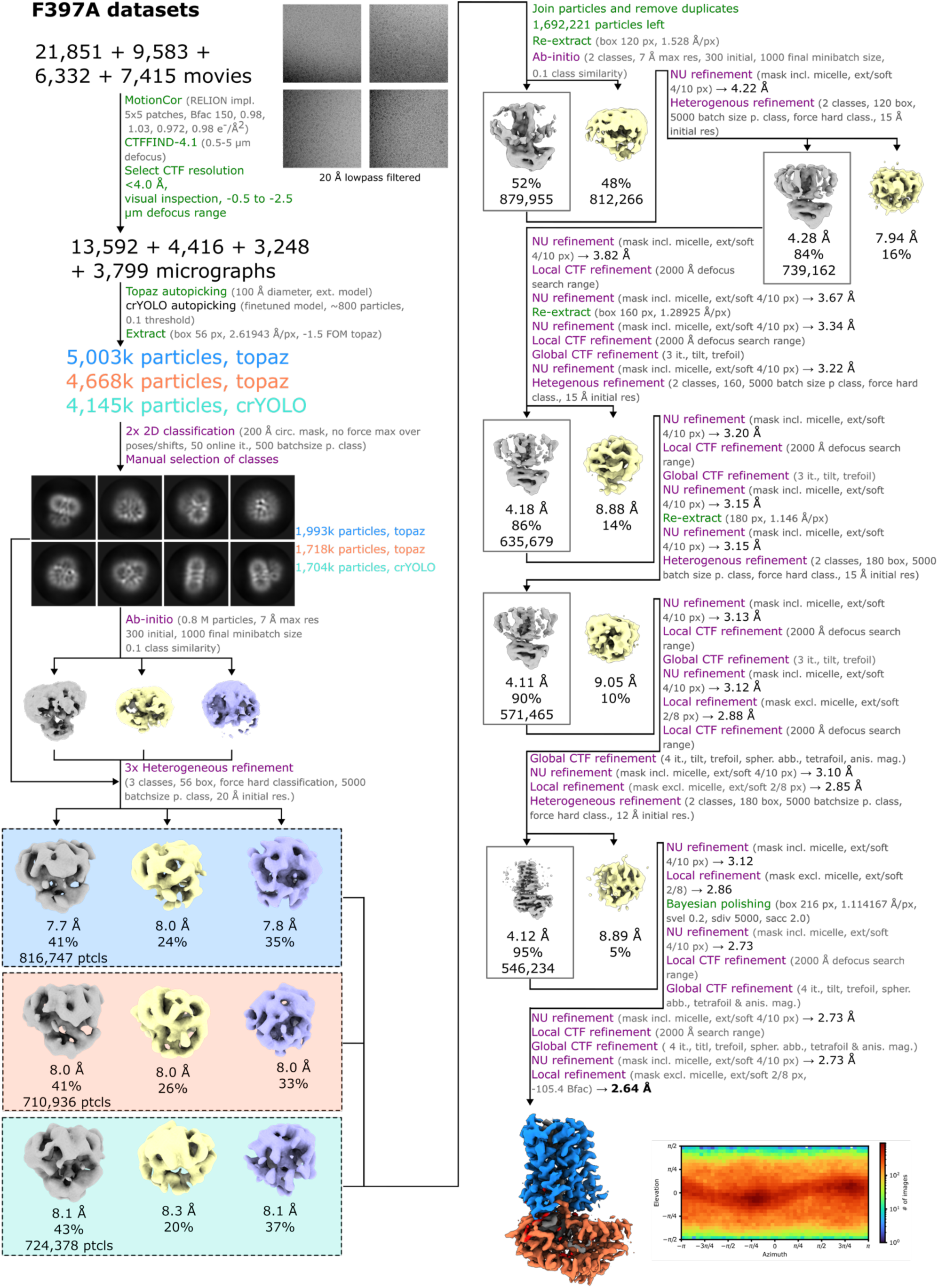
Cryo-EM processing workflow for NADPH-bound Phe397Ala SpNOX reconstruction. Processes run in RELION are colored green, while processes run in cryoSPARC are colored purple. Abbreviations used: External (ext.), pixel (px), per (p.), resolution (res.), particles (ptcls), including (incl.), extension (ext), soft-edge (soft), classification (class.), iterations (it.), excluding (excl.), spherical aberration (spher. abb.), anisotropic magnification (anis. mag.) and B-factor (Bfac).

**Supplementary Fig. 6.**
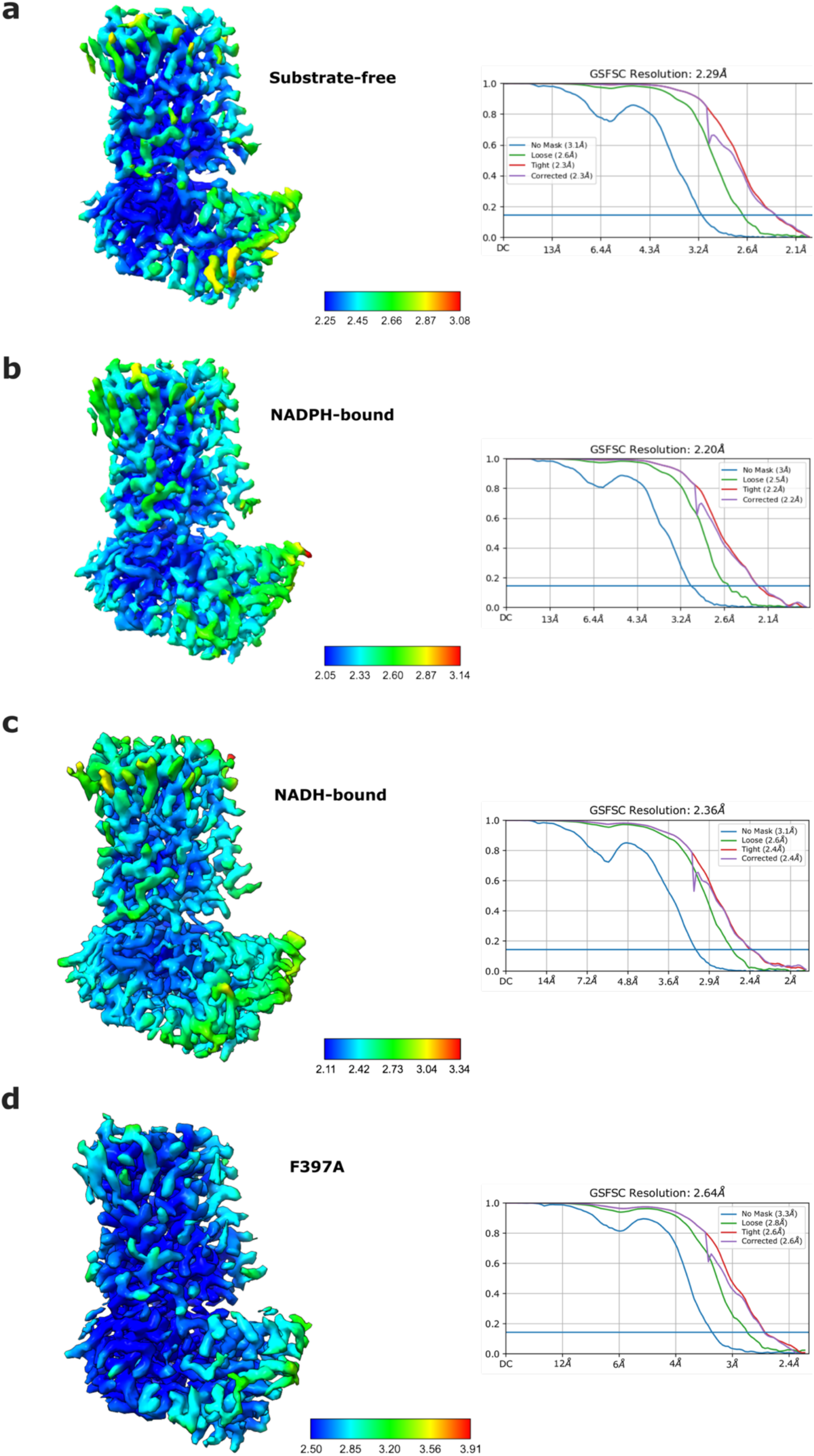
Local resolution and FSC curves of the cryo-EM reconstructions. **a-d,** Left: Density map of the SpNOX reconstruction colored by local resolution. Right: Gold-standard Fourier Shell Correlation (FSC) curve of the 3D reconstructions indicating the resolution at FSC=0.143.

**Supplementary Fig. 7.**
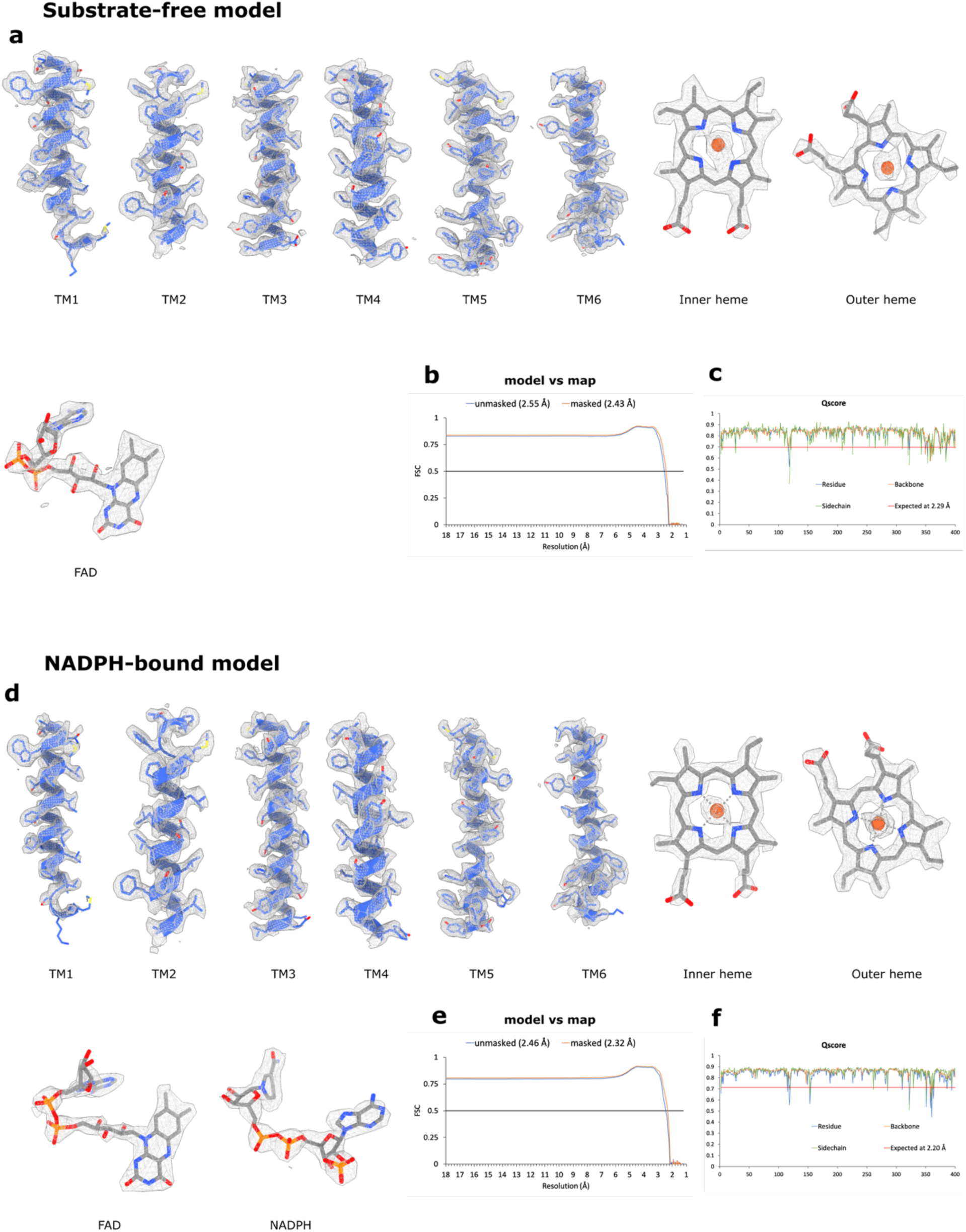
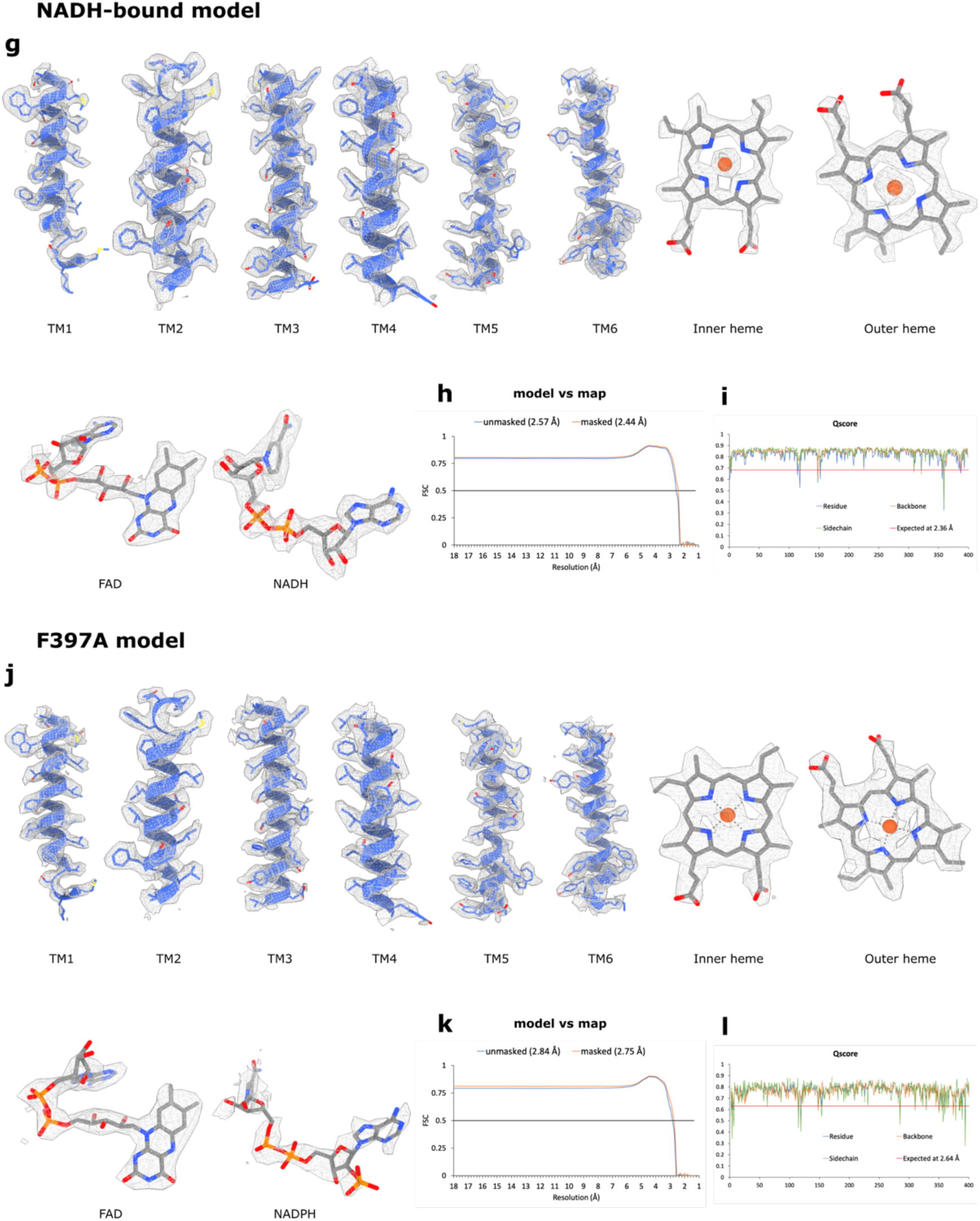
SpNOX model vs map validation. **a,** Density fit of the transmembrane helices and cofactors of the substrate-free model. **b,** Model vs map FSC curve. **c,** Q-score of the residue average, backbone, side chain and expected value for map resolution. **d,** Density fit of the transmembrane helices and cofactors of the NADPH-bound model. **e,** Model vs map FSC curve. **f,** Q-score of the residue average, backbone, side chain and expected value for map resolution. **g,** Density fit of the transmembrane helices and cofactors of the NADH-bound model. **h,** Model vs map FSC curve. **i,** Q-score of the residue average, backbone, side chain and expected value for map resolution. **j,** Density fit of the transmembrane helices and cofactors of the Phe397Ala model. **k,** Model vs map FSC curve. **l,** Q-score of the residue average, backbone, side chain and expected value for map resolution.

**Supplementary Fig. 8.**
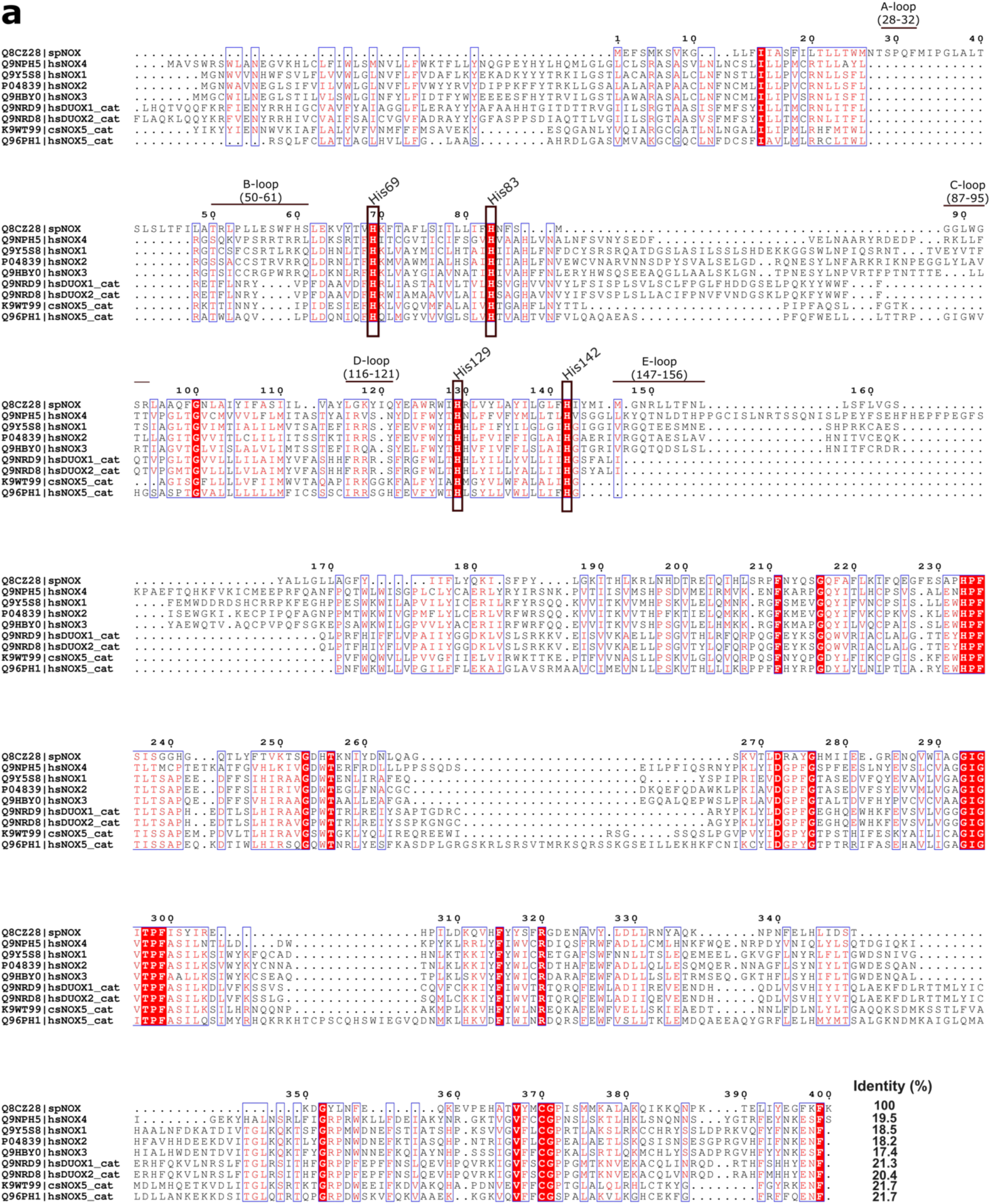
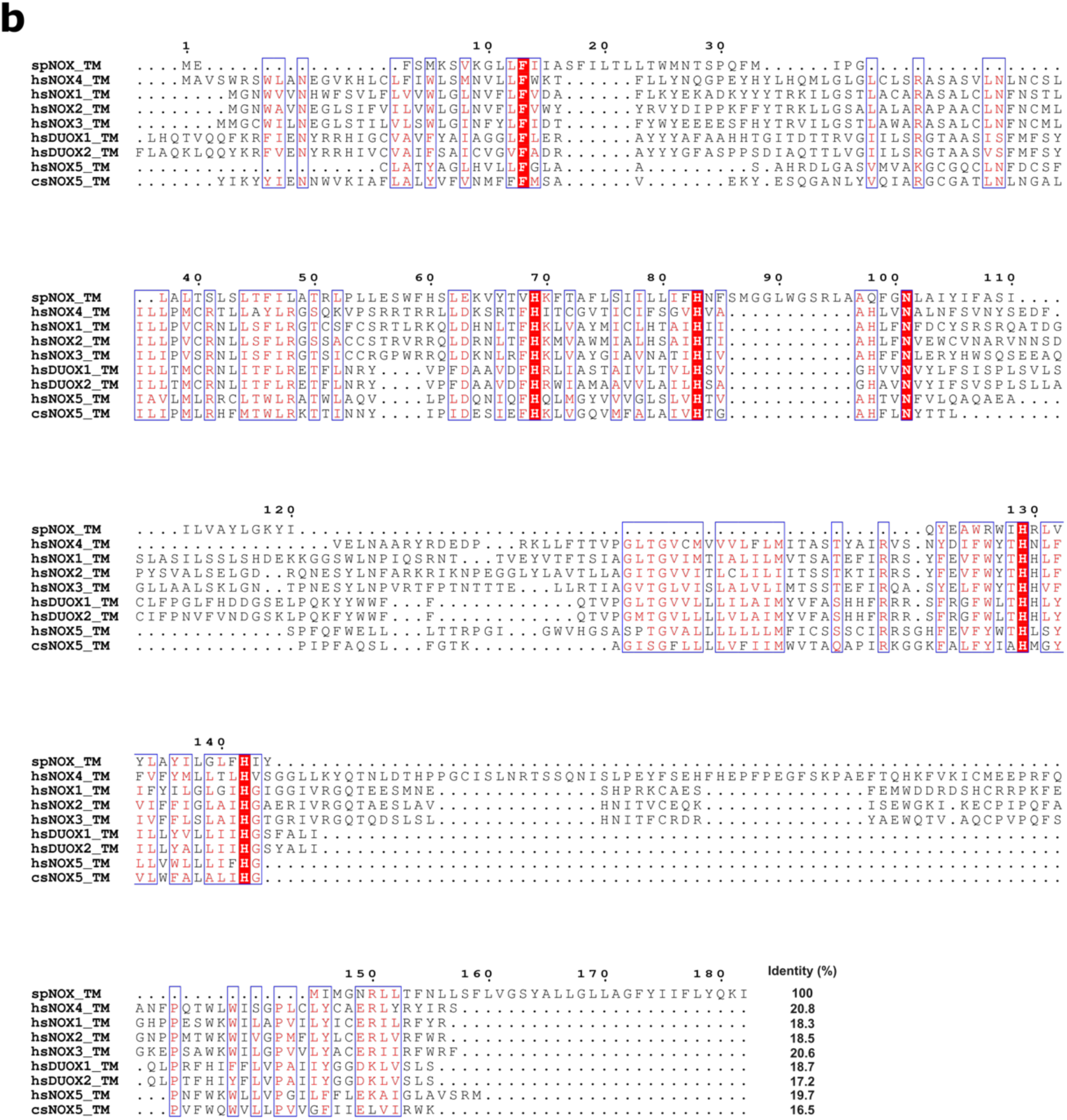
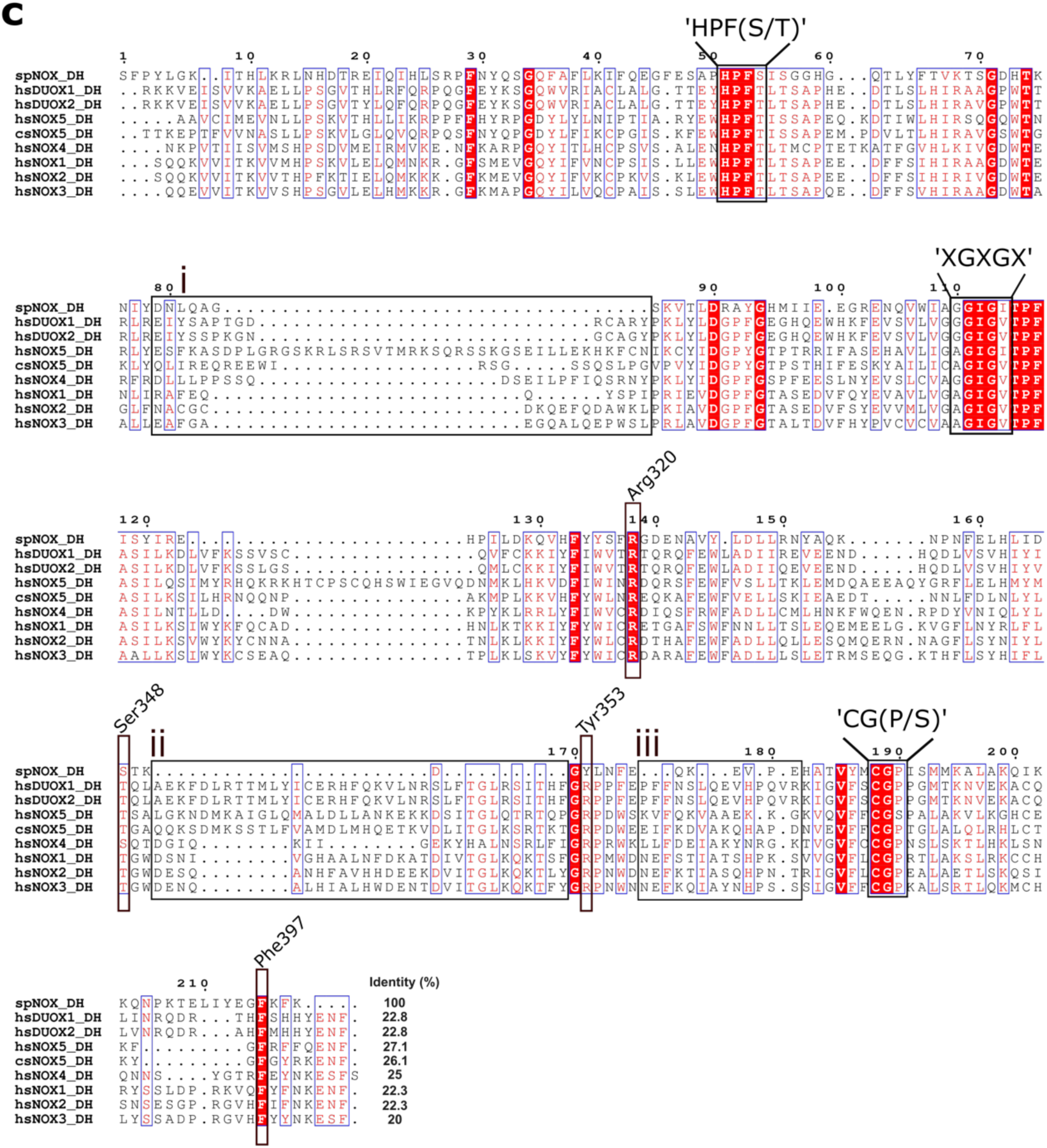
Sequence alignment of SpNOX with human and cyanobacterial NOXs. All sequence identity values were calculated with respect to SpNOX. **a,** Sequence alignment of full-length SpNOX (Q8CZ28) with the catalytic subunits of human (*Homo sapiens*, hs) NOX 1 (Q9Y5S8), NOX2 (P04839), NOX3 (Q9HBY0) and NOX4 (Q9NPH5); and the catalytic core of human NOX5 (Q96PH1, 236-765), human DUOX1 (Q9NRD9, 1027-1551), human DUOX2 (Q9NRD8, 1023-1548) and *Cylindrospermum stagnale* csNOX5 (K9WT99, 210-693). Heme-coordinating histidines (His69, His83, His129 and His142) and the amino acids of the extracellular (A, C and E) and intracellular (B and D) loops of SpNOX are highlighted. **b**, Sequence alignment of SpNOX TM domain (1-182) with the TM domains of human NOX1 (1-290), NOX2 (1-290), NOX3 (1-289), NOX4 (1-305), NOX5 (236-443), DUOX1 (1027-1269), DUOX2 (1023-1266) and csNOX5 (210-409). **c**, Sequence alignment of SpNOX DH domain (183-400) with the DH domains of human NOX1 (291-564), NOX2 (291-570), NOX3 (290-568), NOX4 (306-578), NOX5 (444-765), DUOX1 (1270-1551), DUOX2 (1268-1548) and csNOX5 (410-693). The additional eukaryotic NOX sequences at the DH domain (i, ii and iii as displayed in Supplementary Fig. 9) are boxed; consensus sequences ‘HPF(S/T)’, ‘XGXGX’ and ‘CG(P/S)’, regulatory Phe397 and the amino acids involved in NAD(P)H-binding Arg320, Ser348 and Tyr353 are highlighted.

**Supplementary Fig. 9.**
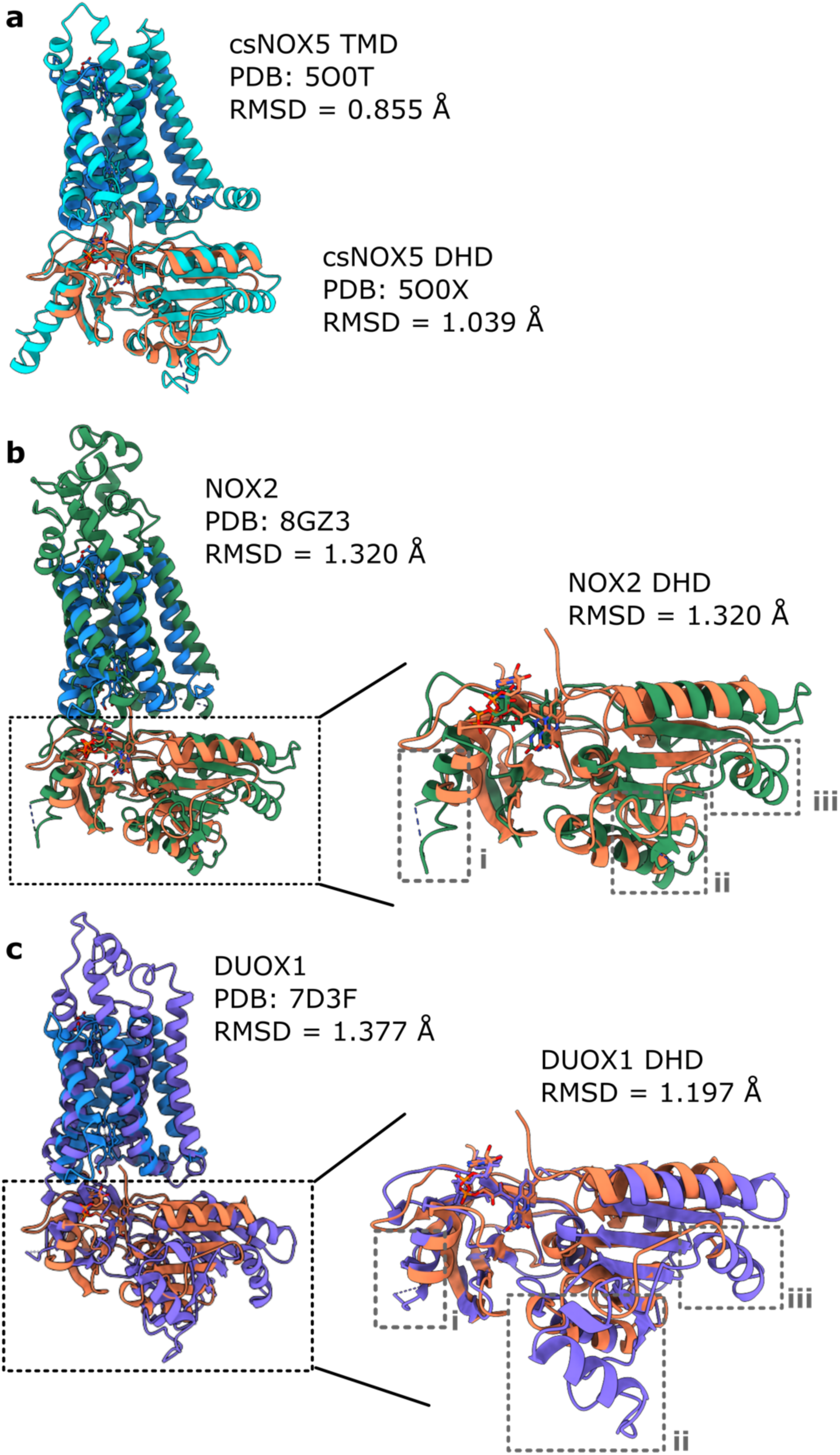
Structural comparison of SpNOX with human and cyanobacterial NOXs. **a,** Structure alignment of the csNOX5 TM (PDB: 5O0T) and DH (PDB: 5O0X)^1^ domains with SpNOX. RMSD values are given for 40 and 131 residue pairs for the TM and DH domains, respectively. **b,** Structure alignment of full-length NOX2 (PDB: 8GZ3)^2^ and DH domain of NOX2 with SpNOX. RMSD values are given for 87 and 78 residue pairs for the full length and DH domain alignments, respectively. **c,** Structure alignment of the catalytic core of human DUOX1 (PDB: 7D3F) ^3^ and DH domain of human DUOX1 with SpNOX. RMSD values are given for 39 and 88 residue pairs for the full length and DH domain alignments, respectively. Additional structural elements absent in SpNOX but present in NOX2 or DUOX1 are indicated by grey boxes.

**Supplementary Fig. 10.**
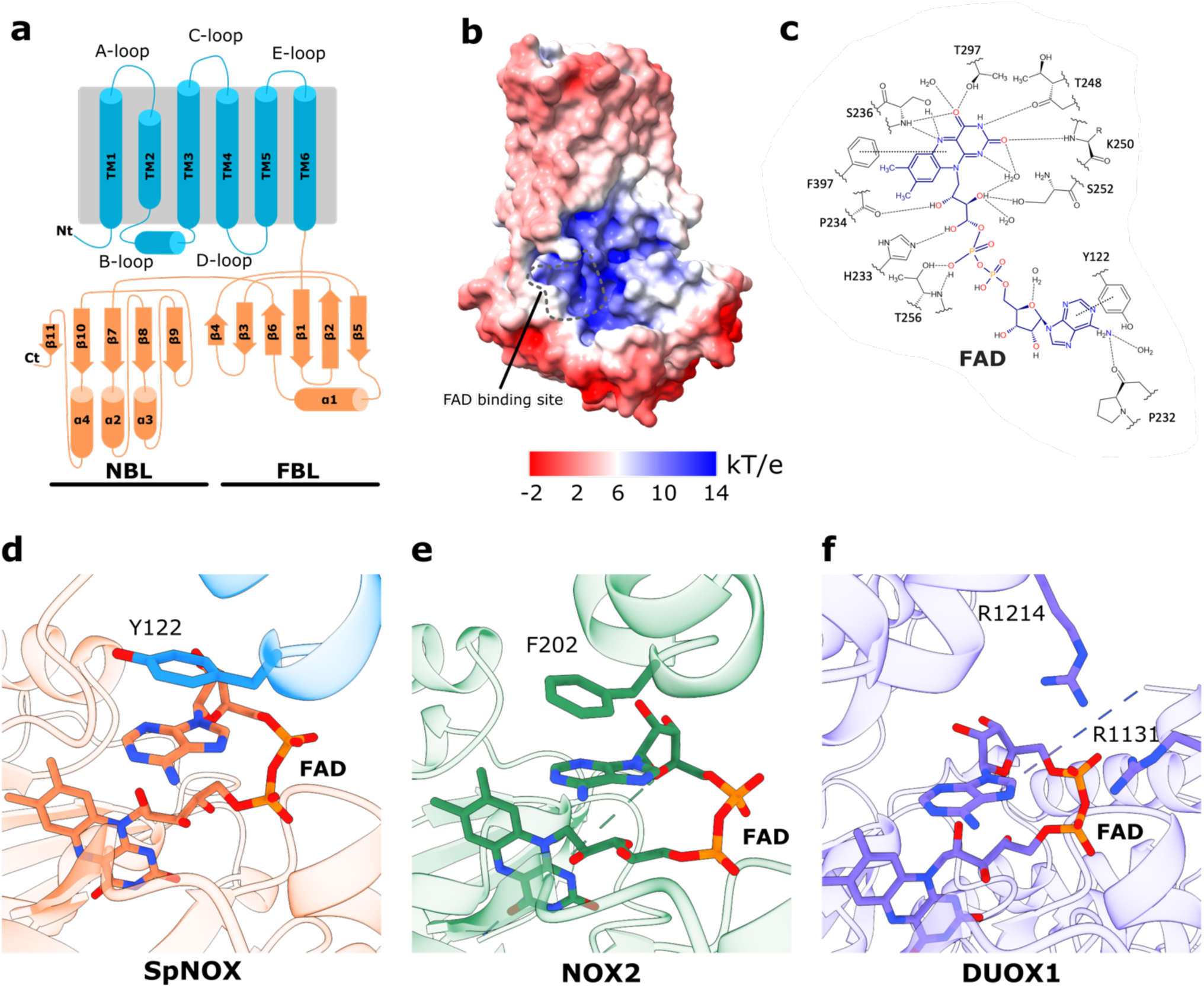
Structural organization of SpNOX and FAD binding. **a,** Cartoon topology model of SpNOX with the lipid bilayer indicated in grey. **b,** Volume representation of SpNOX showing the electrostatic potential with the positively charged FAD-binding site indicated. k, Boltzmann constant; T, temperature (K); e, charge units. **c,** Schematic representation of FAD binding in SpNOX. **d-f,** The geometry of FAD in SpNOX (**d**), NOX2 (PDB: 8GZ3)^2^ (**e**) and human DUOX1 (PDB: 7D3F)^4^ (**f**) is conserved, but in DUOX1 it involves positively charged amino acids at the TM domain (Arg1131 and Arg1214) instead of an aromatic residue (Tyr122 in SpNOX; Phe202 in NOX2).

**Supplementary Fig. 11.**
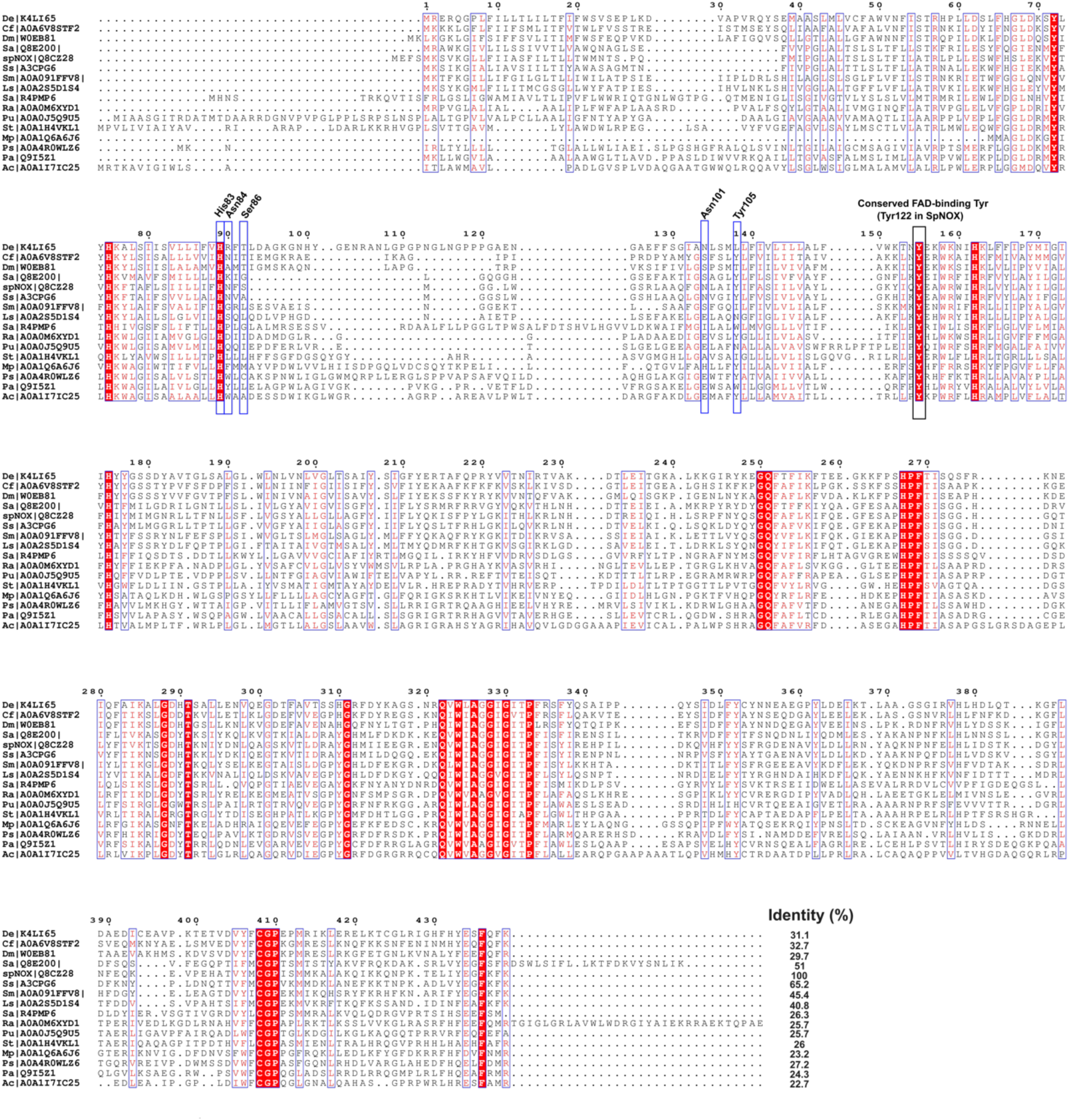
Sequence alignment of bacterial SpNOX homologues. The indicated sequence identity is calculated with respect to the spNOX sequence. The amino acids of the putative reaction centers are boxed in blue. The conserved Tyr for binding FAD (Tyr122 in SpNOX) is boxed in black. **De,** *Dehalobacter sp.*; **Cf,** *Clostridium fungisolvens*; **Dm,** *Desulfitobacterium metallireducens*; **Sa,** *Streptococcus agalactiae*; **spNOX,** *Streptococus penumoniae* NOX; **Ss,** *Streptococcus sanguinis*; **Sm,** *Smithella sp.*; **Ls,** *Lysinibacillus sphaericus*; **Sa,** *Saccharimonas aalborgensis*; **Ra,** *Roseibium aggregatum*; **Pu,** *Puniceibacterium sp.*; **St,** *Streptomyces sp.*; **Mp,** *Mucilaginibacter polytrichastri*; **Ps,** *Paraburkholderia strydomiana*; **Pa,** *Pseudomonas aeruginosa*; **Ac,** *Acidovorans caeni*.

**Supplementary Fig. 12.**
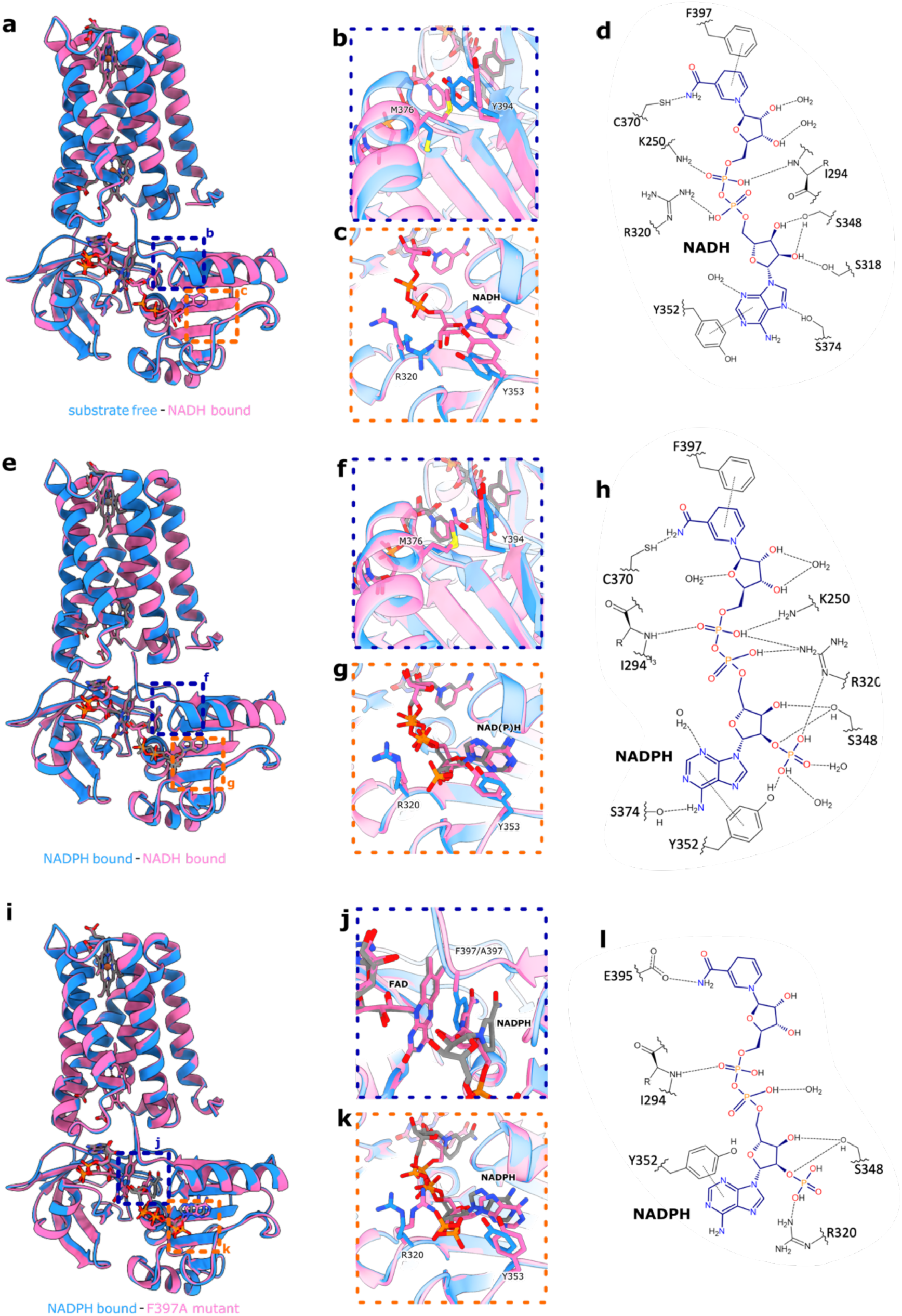
Comparison between the SpNOX structures. **a,** Overlayed cartoon models of substrate-free and NADH-bound SpNOX. **b & c,** A closer look at side chain rearrangements upon substrate binding. **d,** Schematic representation of NADH binding. **e,** Overlayed cartoon models of NADPH-bound and NADH-bound SpNOX. **f & g,** A closer look at side chain rearrangements upon substrate binding. **h,** Schematic representation of NADPH binding. **i,** Overlayed cartoon models of NADPH-bound and F397A SpNOX. **j & k,** A closer look at side chain rearrangements upon substrate binding. **l,** Schematic representation of NADPH binding.

**Supplementary Fig. 13.**
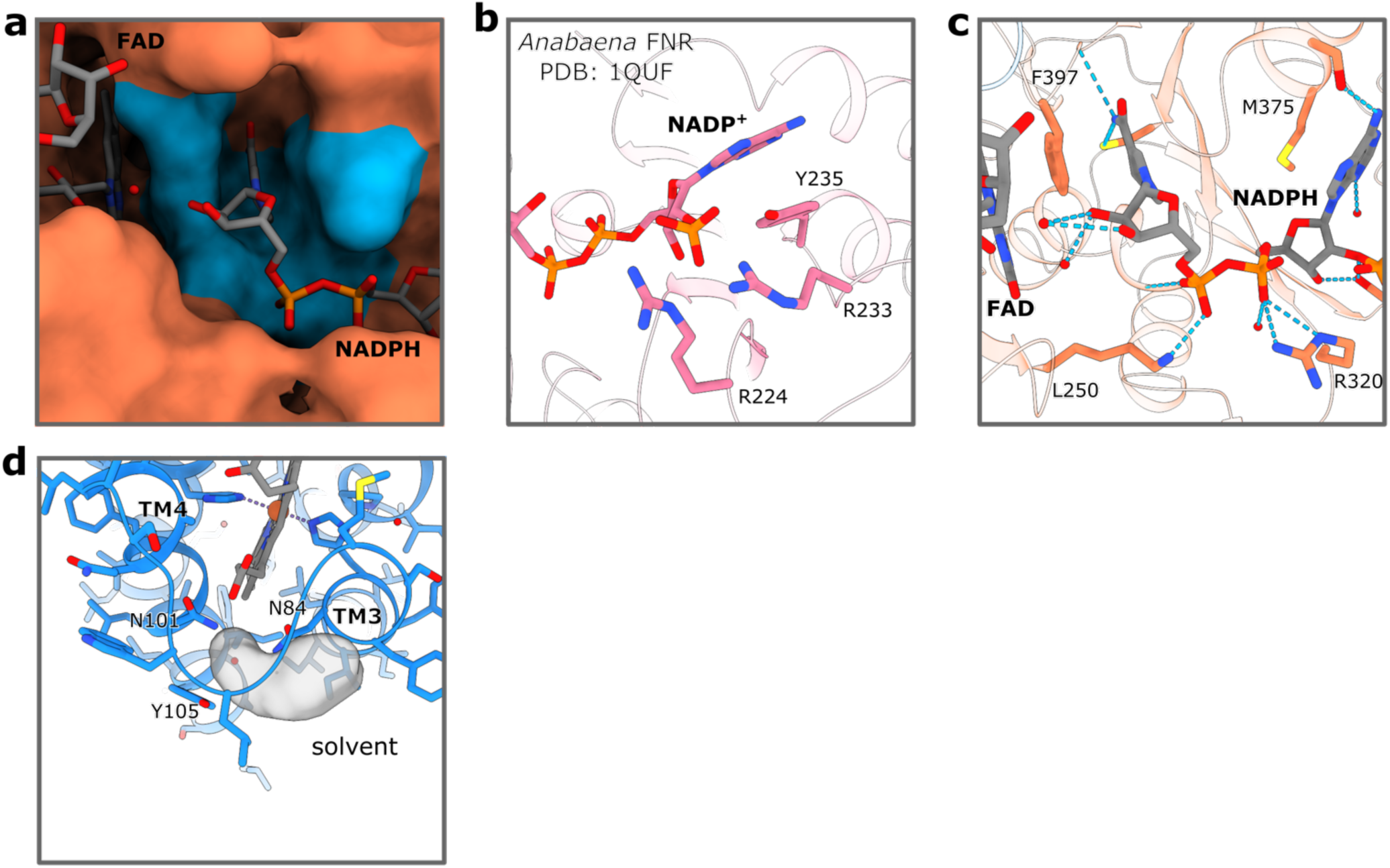
A detailed view of substrate binding in SpNOX. **a,** Surface representation of SpNOX with NADPH bound showing the cavity to accommodate nicotinamide formed by the amino acids of the consensus sequences ‘XGXGX’ and ‘CG(S/P)’, and by Phe397 (all colored cyan). **b,** Substrate binding of *Anabaena* FNR (PDB: 1QUF) ^5^ showing the polar interactions between the protein and the 2’-phosphate via arginine residues. **c,** A closer look at the nicotinamide-bound ribose of NADPH reveals the absence of direct interactions with SpNOX. **d,** A potential O_2_/O_2_^•−^ entrance and exit path at one of the proposed oxygen binding sites was mapped using the Hollow tool.

**Supplementary Fig. 14.**
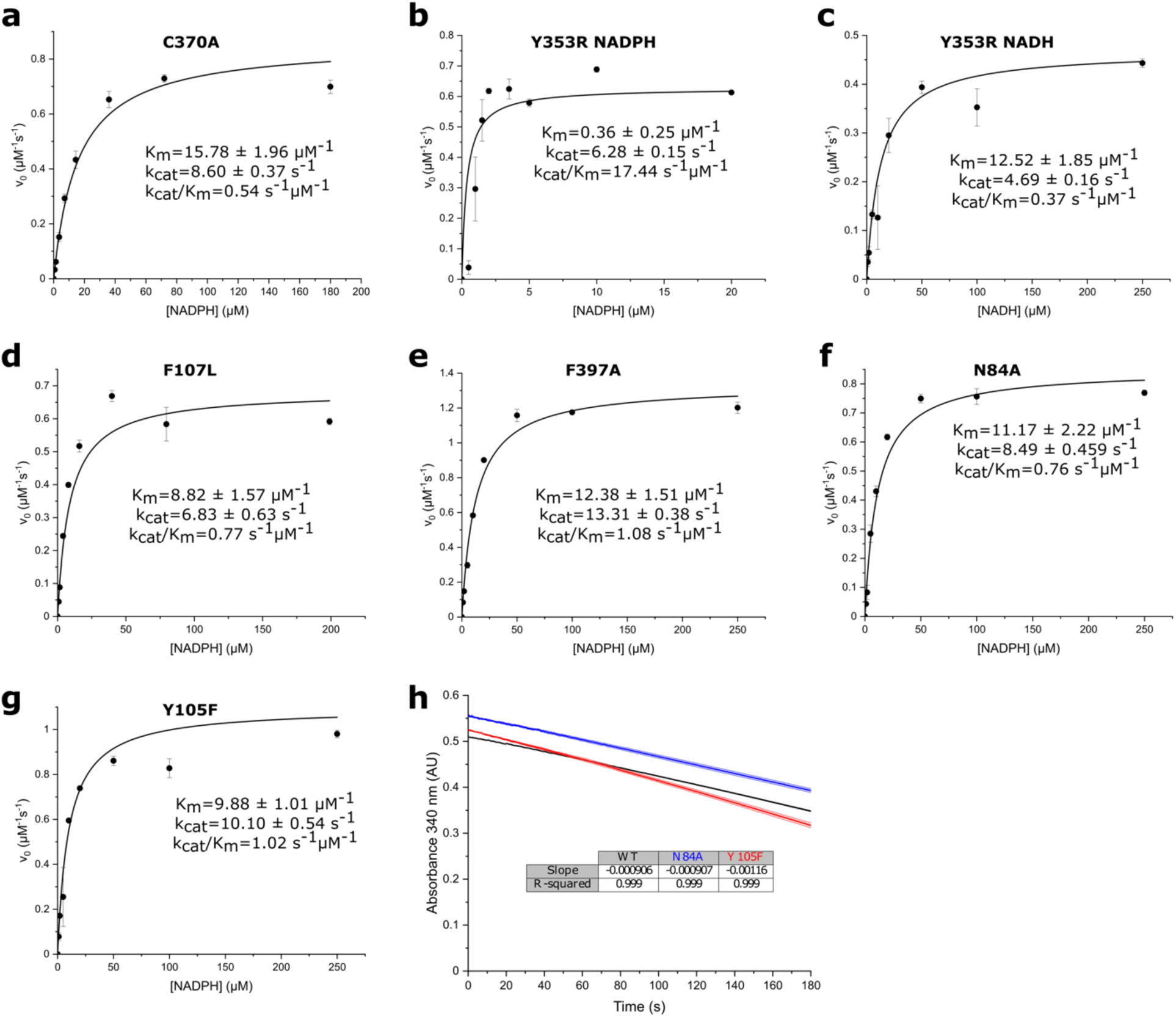
The NAD(P)H-oxidase activity of SpNOX mutants under steady state conditions. **a-g,** Data obtained from a cytochrome c reduction assay was fitted to the Michaelis-Menten equation to obtain apparent K_m_ and k_cat_ values. **h,** Data obtained from NADPH oxidation assay. A linear function was fit to obtain the slopes. The NADPH oxidation rates (µM s^-1^) are calculated using an NADPH extinction coefficient of 6.22 L mmol^-1^ cm^-1^. Mean values of three technical replicates are plotted and SEM is indicated. Data for individual replicates are available in the source data.

**Supplementary Fig. 15.**
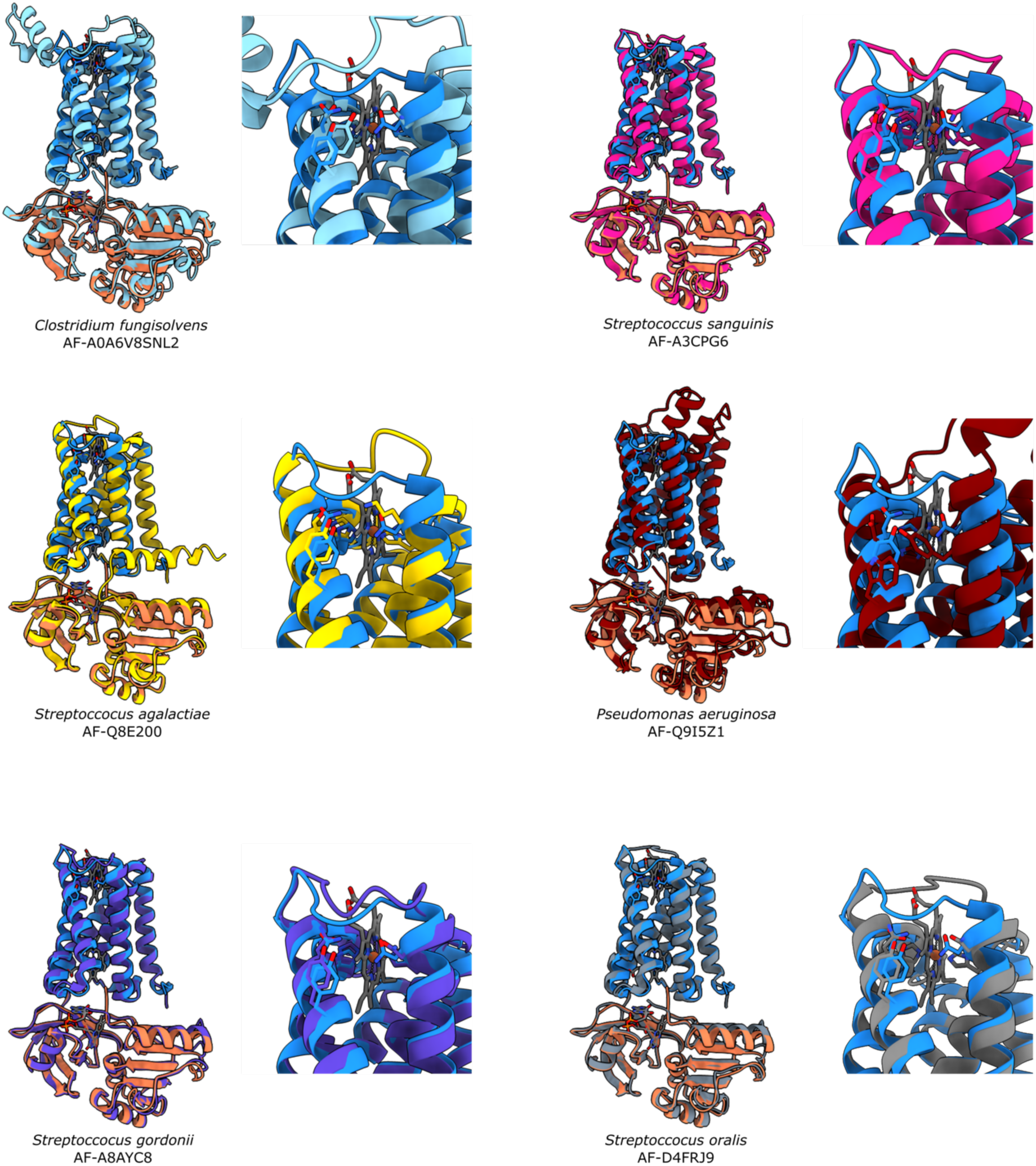
Predicted structures of bacterial SpNOX-like protein generated by AlphaFold2 showing residue conservation near the outer heme, highlighted in supplementary figure 12.

**Supplementary Fig. 16.**
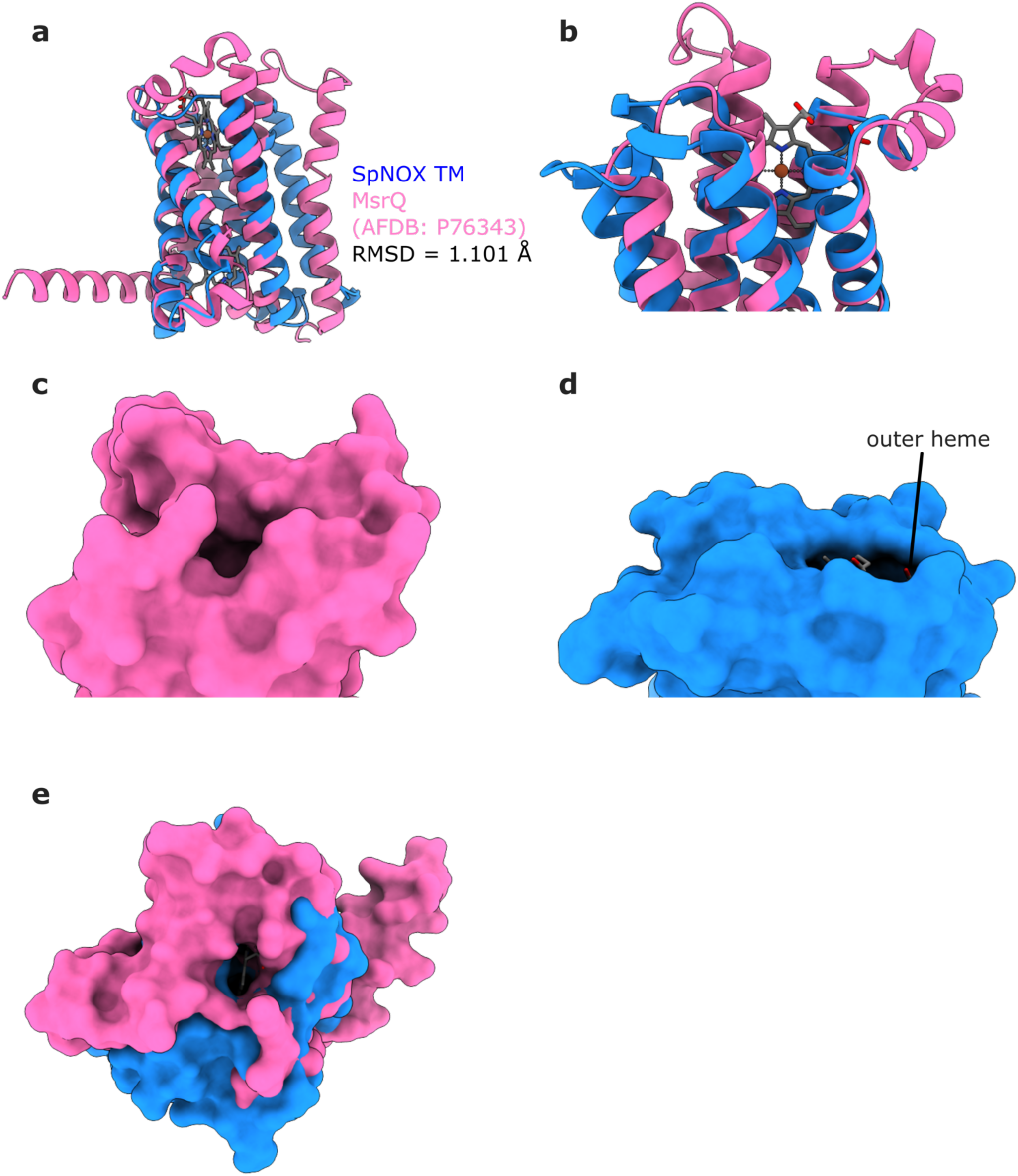
SpNOX shows an exposed outer heme similar to its homologue MsrQ. **a,** Cartoon representation of the AlphaFold model of MsrQ (AlphaFold DB: P76343) aligned to SpNOX TMD. RMSD value is given for 38 residue pairs. **b,** A closer look at the periplasmic region of SpNOX and MsrQ. **c,** Surface representation of MsrQ extracellular region showing a large cavity. **d,** Surface representation of SpNOX periplasmic region showing the solvent-accessible outer heme. **e,** Top view of the overlayed surface representations of SpNOX and MsrQ.

**Supplementary Table 1.**
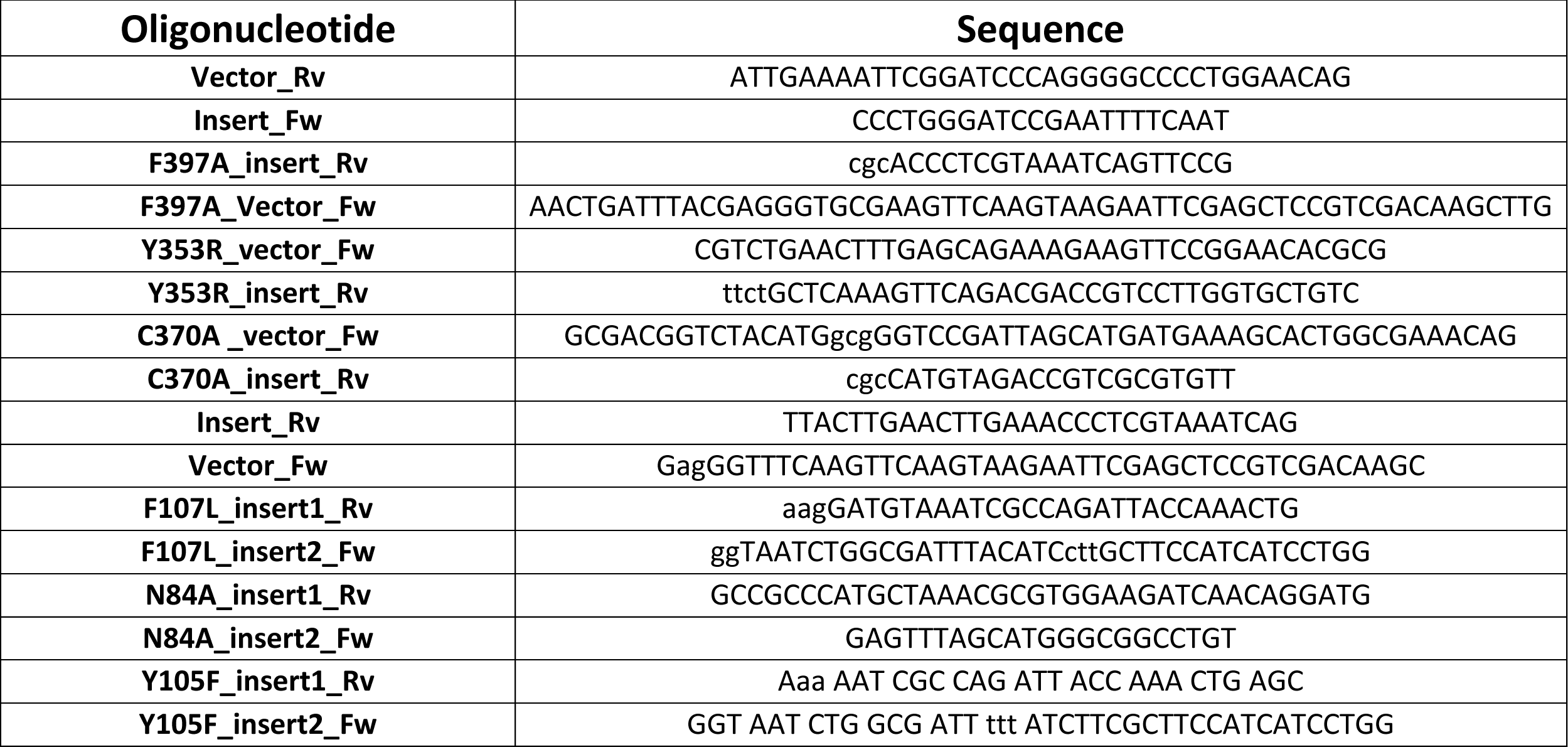
Oligonucleotides used to obtain the fragments for the NEBuilder Hifi Assembly reactions to produce the mutants.

**Supplementary Table 2.** Oligonucleotide combinations used in PCR to obtain the DNA fragments for the NEBuilder HiFi DNA Assembly reactions to generate SpNOX mutants. A pET28a vector with WT SpNOX between NcoI and EcoRI sites was used as template in all reactions.

| Mutant | Insert1 |  | Insert2 |  | Linearized vector |  |
| --- | --- | --- | --- | --- | --- | --- |
|  | Insert1_Fw | Insert1_Rv | Insert2_Fw | Insert2_Rv | Vector_Fw | Vector_Rv |
| F397A | Insert_Fw | F397A_insert_Rv | - | - | F397A_Vector_Fw | Vector_Rv |
| Y353R | Insert_Fw | Y353R_insert_Rv | - | - | Y353R_vector_Fw | Vector_Rv |
| C370A | Insert_Fw | C370A_insert_Rv | - | - | C370A_vector_Fw | Vector_Rv |
| F107L | Insert_Fw | F107A_insert1_Rv | F107L_insert2_Fw | Insert_Rv | Vector_Fw | Vector_Rv |
| N84A | Insert_Fw | N84A_insert1_Rv | N84A_insert2_Fw | Insert_Rv | Vector_Fw | Vector_Rv |
| Y105F | Insert_Fw | Y105F_insert1_Rv | Y105F_insert2_Fw | Insert_Rv | Vector_Fw | Vector_Rv |

